# OmicGlaze: Spatial Multi-Omic Mapping of Traumatic Brain Injury

**DOI:** 10.64898/2026.03.30.715462

**Authors:** Yiheng Li, Sam J. Neuffer, Joseph Wider, Sai Ma, Neil Zhao, Liam McCracken, Thomas Sanderson, Jing-fei Dong, Yanxiang Deng, Yang Xiao

## Abstract

Traumatic brain injury (TBI) is a major cause of mortality and long-term disability worldwide, giving rise to complex neurological complications that impact millions of individuals each year. Cellular stress and neuronal injury vary dramatically across cortical layers, vascular niches, and between the ipsilateral (injured) or contralateral (uninjured) hemispheres. There is a critical need for quantitative measures that capture the spatial distribution of injury-induced cellular changes, as well as the gene regulatory elements that drive them. Here, we developed *OmicGlaze*, an experimental and computational workflow for systematically profiling the spatial transcriptome and epigenome of mouse brains following mild traumatic brain injury. We established a spatial scoring system, and identified region-specific biological processes post injury, including changes in neuronal activities, cellular stress, immune response, and gliosis. Spatial assay for transposase-accessible chromatin with sequencing (Spatial ATAC-seq) generated the first epigenetic map of traumatic brain injury near single-cell resolution. Notably, we identified the Activator Protein-1 family transcription factor Atf3 as a key gene regulator of injury-induced cellular stress. Together, these spatial multi-omics analyses revealed gene regulatory network in TBI and provided a broadly applicable framework for dissecting cellular and molecular mechanisms underlying complex neurological disorders.

## Introduction

Traumatic brain injury (TBI) is a major public health concern that affects approximately 64 million individuals worldwide, including athletes, veterans, and survivors of accidents, falls, and interpersonal violence [1, 2]. Brain damage arises from both the initial insult and subsequent secondary injuries including uncontrolled hemorrhage, cerebral edema, focal or diffuse axonal injury, tissue ischemia, and cerebral and systemic inflammation [3–5]. TBI carries profound long-term consequences for cognitive, emotional, and physical health, imposing substantial burdens on patients, their families, and the healthcare system [6]. However, mechanisms that drive the progression of a localized injury into widespread damage across distinct brain regions over subsequent days remain unclear. In addition, although the type of these secondary injuries is well-documented, their cellular and molecular characteristics are poorly defined within the tissue context, making it challenging to predict the time and location of secondary injuries and identify targeted therapeutic intervention.

Recent advances in single-cell RNA sequencing (scRNA-seq) technologies have enabled the interrogation of heterogeneity in the injured brain by systematically evaluating diverse cell populations at large-scale [7–10]. For instance, single-cell RNA-seq profiling of over 330,000 cells at 24 hours and 6 months post injury from the mouse peri-contusion cortex was reported, revealed cell-type-specific immune response in the secondary damage [11]. However, neurons cell population was under-represented in this work (less than 3%) [11], which typically make up 46% of the mouse brain cell population [12]. This skewed representation could lead to a biased analysis into neuronal injuries expression changes after TBI. Furthermore, dissociating brain tissue into individual cells lose the spatial information, preventing region-specific studies and neighborhood analyses of injury-induced changes, as well as signals from subcellular compartments, such as axonal and dendritic processes [13]. Understanding region-specific changes is critical in therapeutic intervention, as it reveals where and how cells alter their states and function in response to injury. For example, microglia exhibit diverse cellular function at different brain regions. Homeostatic microglia disperse throughout the brain, as a sentinel to monitor the microenvironment [14], whereas activated microglia localize to injured neurons or vasculature, releasing cytokines and phagocytosing damaged cells [15].

Two spatial mRNA-seq studies have begun addressing these gaps. Using the Visium V1 platform (10X Genomics), Swaro *et al*. profiled the transcriptomic landscape at 7 days post TBI [16], and Kounelis-Wuillaume *et al*. examined dynamic gene expression signatures at both 2 and 7 days post-injury, both highlighted the immune cell activation in injury responses [16, 17]. Both studies mapped only the injured hemisphere, due to a limited capture area (6.5 mm x 6.5 mm). As a result, neither examines changes in the contralateral versus ipsilateral hemispheres, a comparison that is essential for accurately assessing injury-induced alterations within the same individual. In addition, peri-injury differences across distinct coronal planes were not evaluated. Even within the same brain, cellular damage varies depending on the distance from the injured site. Without mapping the contralateral hemisphere and sampling across peri-injury coronal planes, these analyses offered an incomplete assessment of injury-induced changes.

Moreover, these studies focused on individual differentially expressed genes without integrative approaches to resolve the spatial neighborhoods or tissue-level pathway organization. They analyzed the spatial data as single cell RNA or pseudo-bulk RNA data, without evaluating the continuous changes in each spatial anatomical region. Most importantly, spatial transcriptomics alone measures gene expression but cannot capture the epigenetic regulatory elements that control transcription. Fully understanding the mechanisms underlying neuronal damage and post-injury remodeling therefore require new technologies capable of profiling non-coding regions of the epigenome.

To overcome these limitations, we have developed *OmicGlaze*, a sequencing-based experimental workflow optimized for TBI and a computational pipeline that investigates both gene expression and gene regulation with spatial contexts. *OmicGlaze* consists of two state-of-art technologies: spatial mRNA-seq for gene expression mapping and spatial ATAC-seq [18] for chromatin accessibility profiling, both at centimeter-scale tissue coverage. By profiling two modalities *in situ* without tissue dissociation, our spatial assays preserve both spatial information and subcellular features, including axonal and dendritic processes. *OmicGlaze* enables a comprehensive characterization of the dynamic, heterogeneous, and spatially organized responses of the brain after TBI. It enhances resolution in spatial biology and enables high throughput readouts of cell states, much like each glaze in a painting shaping the final visual.

## Results

### Spatial mapping of the whole transcriptome in TBI brains

To comprehensively characterize the changes induced by a lateral impact-induced mild traumatic brain injury, we developed a tri-omics workflow including histology, transcriptomics, and epigenetics (Fig. 1A). The incorporation of these three omics modalities is essential, as each provides complementary information. Conventional histological methods such as Hematoxylin and Eosin (H&E) or Nissl staining provide morphological insights but do not capture molecular-level functional changes. For example, in a Nissl-stained section of a brain with moderate TBI introduced to the top right hemisphere, it was challenging to assess injury severity or identify significant morphological changes, as tissue integrity remained largely intact (Fig. 1B). From the zoom in view of the two sides highlights the cortex region, the cell density and morphology were not disrupted by the mild injury (Fig. 1C, D). To validate this, we calculated the cell number using CellPose-SAM algorithm [19], based on nuclei in the adjacent H&E images of the TBI samples. Every nucleus was counted and assigned to a hemisphere with a spatial mask, and enumerated to obtain cell counts for the uninjured (left) and injured (right) sides (Additional file 1: Fig. S1A). The analysis confirmed that the total number of cells were not significantly different between uninjured and injured hemisphere (N1 = 5, N2 = 5, p-value = 0.79). Traditional histology cannot detect functional changes in mild traumatic brain.

**Figure 1.**
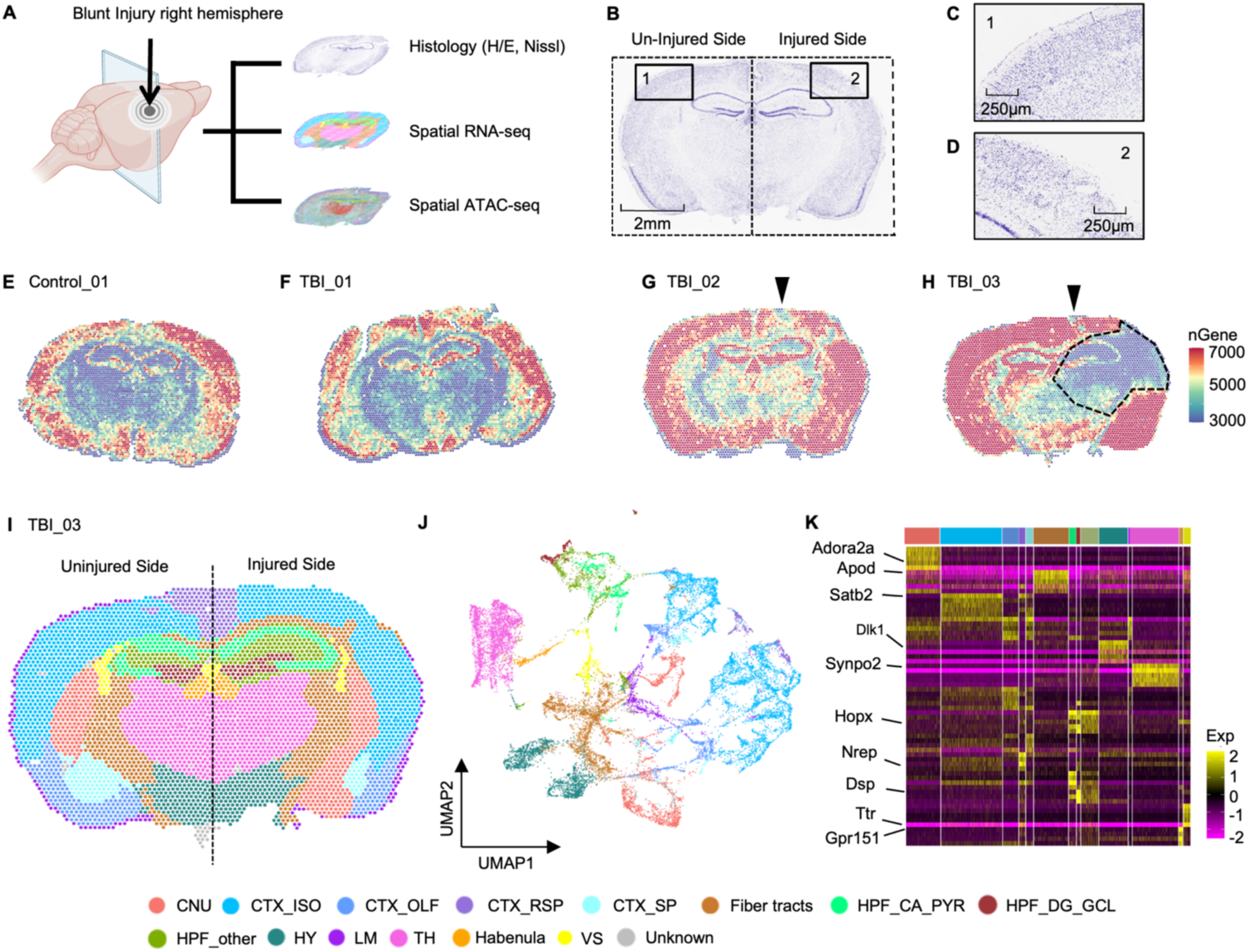
Study design and spatial annotation of mouse brain regions. **A** A focal blunt injury was induced in the upper right hemisphere of the mouse brain. Serial coronal sections were obtained at the mid-hippocampal level, with adjacent sections allocated for histological analysis (hematoxylin and eosin staining and Nissl staining), spatial RNA sequencing (spatial RNA-seq), and spatial ATAC sequencing (spatial ATAC-seq). **B** Image of a Nissl-stained section adjacent to the TBI_03 sample used for sequencing, illustrating tissue morphology. The left side corresponds to the uninjured hemisphere, and the right side to the injured hemisphere. **C** Zoomed-in view of the isocortex on the uninjured side of panel B. Scale bar represents 250 µm. **D** Zoomed-in view of the isocortex on the injured side of panel B. Scale bar represents 250 µm. **E** Spatial distribution of the number of detected genes per spot in the Control_01 sample. Each dot represents a 55-micron spot on the Visium slide (11 x 11 mm^2^). **F - H** Spatial distribution of the number of detected genes per spot in the TBI_01, TBI_02, and TBI_03 samples. **I** Spatial region annotation of the TBI_03 sample, highlighting 14 anatomical regions. Abbreviations: CNU, cerebral nuclei; CTX_ISO, isocortex; CTX_OLF, olfactory cortex; CTX_RSP, retrosplenial area; CTX_SP, cortical subplate; HPF_CA_PYR, hippocampal formation cornu ammonis pyramidal layer; HPF_DG_GCL, hippocampal formation dentate gyrus granule cell layer; HPF_other, the polymorphic layer and molecular layer of hippocampus; HY, hypothalamus; LM, leptomeninges; TH, thalamus; VS, ventricular systems. **J** Unsupervised clustering of the integrated dataset comprising Control_01, TBI_01, TBI_02, and TBI_03 samples, colored by brain regions as defined in panel I. **K** Heatmap showing the expression levels of the top five marker genes for each of the 14 annotated regions in Control_01. Each row represents one gene and each column under the same color represents one region. Selected marker genes are labeled on the plot: *Adora2a* (marker for CNU region), *Apod* (fiber tracts), *Satb2* (CTX_ISO region), *Dlk1* (HY region), *Synop2* (TH region), *Hopx* (*HPF_other* region), *Nrep* (CTX_RSP region), *Ttr* (VS region), and *Gpr151* (habenula region).

In this study, we offered transcriptomic profiles as functional readouts post TBI, from one control brain (Control_01) and three TBI samples (TBI_01, TBI_02, and TBI_03). TBI_01 was collected from an independent biological sample 3 hours post TBI. TBI_02 and TBI_03 were collected from a same mouse 24 hours post TBI. TBI_02 represented the caudal plane of hippocampus, as a peri-injury coronal plane, while TBI_03 is 300 μm rostral to the coronal plane of TBI_02, as the center plane near the injury site. In all TBI samples, focal blunt injury was applied to the right hemisphere of the mouse. We evaluated the spatial distribution of the number of genes and the number of transcripts (unique molecular identifiers, UMIs) detected per spot across all four samples. These quality check revealed well-defined tissue boundaries with high gene capture across tissue sections (Fig. 1E-H, Additional file 1: Fig. S1B-G, Table 2). On average, we detected 5,274 ± 683 genes and 14,872 ± 4,169 UMIs per spot across the four samples. The observed variation in gene counts followed expected anatomical structures rather than technical artifacts. These results indicated that the data were of high quality and suitable for downstream spatial transcriptomic analyses. Interestingly, analysis of global gene expression revealed spatial heterogeneity in total gene counts among different anatomical regions of a sample, and also the same anatomical region between control and TBI samples. In the Control_01 sample, around 6000 to 7000 genes were detected in the cortex, striatum, hypothalamus, as well as in the cornu Ammonis and dentate gyrus of the hippocampus. While in other regions (thalamus and fiber tracts), lower number of genes, around 3000 to 5000, was detected (Fig. 1E). TBI_01 also showed similar heterogeneous gene distribution, with around 6000-7000 genes detected at the isocortex, and cornu Ammonis and dentate gyrus of the hippocampus, with low detection in other regions, particularly within the fiber tracts, where about 3,000 genes were detected (Fig. 1F). In contrast, the TBI_03 sample had an distinct pattern: a significant reduction in gene counts to around 3000 in the injured hemisphere, while in other regions: cortex, hypothalamus, cornu Ammonis and dentate gyrus of the hippocampus more than 7000 genes were detected. This decrease was located at the top right region of the isocortex and the upper half of the striatum. This reduction gradually dissipated in radial directions shown in the dashed highlighted region (Fig. 1H). The distribution of the gene number reduction matches the pattern of pressure gradient in lateral TBI [20, 21]. Since the adjacent Nissl-stained slide did not show a significant decrease of cell number and density (Fig. 1B), the decrease in gene counts in the injured hemisphere is due to cell stress not cell death, as measured earlier in Additional file 1: Fig. S1A, no significant difference was observed in cell number counts across two hemispheres. In TBI_02 (peri-injury plane), which is located 300 μm rostral to the injury center, the asymmetric pattern was less pronounced than in TBI_03 (Fig. 1G). The gene distribution overall was symmetric, with more than 7000 gene counts in cortex, hypothalamus, cornu Ammonis and dentate gyrus of the hippocampus except a small area highlighted with an arrow in the figure which is the direct injury site. This pattern suggested that gene expression changes were specific to the injury location.

Our workflow demonstrates that gene expression changes following TBI are highly region-specific, reflecting localized responses to injury that cannot be detectable by conventional histopathology alone. Our method could quantitatively review the entire coronal sections at high resolution (55 μm), allowing comparison in TBI versus control brains, as well as in peri-injury versus injury center planes. The created spatial transcriptomic atlas (>18,000 genes, mapped at 3hr, 24hr post TBI) provided a resource database for the community to check any genes of interest.

### Comprehensive Spatial Annotation of Main Mouse Brain Regions

We next sought to identify the main mouse brain regions regardless of injury type. We initially performed unsupervised clustering in *Seurat* [22], then assigned a region name to each cluster based on the Allen Institute Mouse Brain Atlas [23]. In total, we identified 14 different regions: cerebral nuclei, cortex isocortex, olfactory cortex, cortical subplate, fiber tracts, hippocampal formation cornu Ammonis pyramidal layer, hippocampal formation dentate gyrus granule cell layer*, HPF_other* region which represents the polymorphic layer and molecular layer of hippocampus, hypothalamus, leptomeninges, retrosplenial area, thalamus, habenula, ventricular systems, and one cluster for areas that could not be clearly classified (Fig. 1I). These 14 regions were identified in all four samples (Additional file 1: Fig. 1I-K) and were regions expected based on previous analyses [10, 12, 23].

We integrated the spatial data from all samples using canonical correlation analysis (CCA) [22], followed by UMAP dimensionality reduction. We visualized the integrated data in two ways: a UMAP plot colored by annotated anatomical regions and a UMAP plot colored by sample identity. In the region-based UMAP, each anatomical region formed a distinct cluster (Fig. 1J), supporting the accuracy of our regional annotations. The sample-based UMAP showed that spots from different samples were intermingled within clusters rather than segregating by sample (Additional file 1: Fig. 1H). This pattern indicates that the clustering reflects underlying biological variation rather than technical batch effects. To further confirm the annotated regions, we performed differential gene expression analysis and found that marker genes aligned well with expected expression in annotated regions (Fig. 1K). For example, *Adora2a,* which plays a key role in integrating neural signals to regulate cognition, is highly specific to the cerebral nuclei [24]. *Satb2* is a critical transcription factor for the development and differentiation of upper-layer cortical neurons; its loss in the cerebral cortex induces cellular senescence, supporting its status as a representative marker for the isocortex [25]. *Synpo2*, an actin-binding protein, is predominantly expressed in the thalamus [26]. *Gpr151* is highly expressed in the habenula, where it regulates synaptic plasticity [27]. Each of these genes has strong regional enrichment consistent with in situ hybridization data from the Allen Mouse Brain Atlas [12] (Additional file 1: Fig. S2A-D). We then manually separated the brain further into injured and uninjured sides, assigning this side information to each cluster so that each region on one side has a corresponding counterpart on the other. This high-accuracy regional annotation is critical for downstream analyses, as it enables the identification of region-specific alterations and facilitates differential gene expression comparisons across paired regions.

### Spatially Analysis of Marker Genes of Key Biological Processes

Next, we examined six key cellular processes previously reported to be altered following TBI [28, 29]. For each process, we selected one spatially variable gene and visualized its spatial distribution in the integrated Control_01 and TBI_03 samples using CCA. All six genes showed significant changes in expression levels after TBI, especially on the injury side.

Astrogliosis is the process by which astrocytes become reactive in response to injury, disease, or inflammation. It serves protective functions by isolating damaged tissue, removing debris, and modulating inflammation. However, excessive or chronic astrogliosis can lead to the formation of a glial scar, which may hinder neuronal regeneration and impair recovery [30]. Glial scarring has also been reported to contribute to the development of epilepsy [31]. Notably, astrogliosis is one of the hallmark responses to TBI [28, 29]. *Gfap*, which encodes an intermediate filament protein that is strongly upregulated in activated astrocytes, is a maker for astrogliosis [32]. Compared to the Control_01 sample, the spatial expression of *Gfap* was significantly increased in certain regions of the TBI_03 sample. The increase was considerable in the hippocampus region on the injury side especially the *HPF_other* region, as well as regions like fiber tracts, or leptomeninges (Fig. 2A). The distribution of gene expression levels in spatial spots is significantly higher on the injured side of *HPF_other* region than the same region of the uninjured side (Fig. 2B). Elevated expression of *Gfap* suggests that there is robust astrocyte activation in certain brain regions following injury.

**Figure 2.**
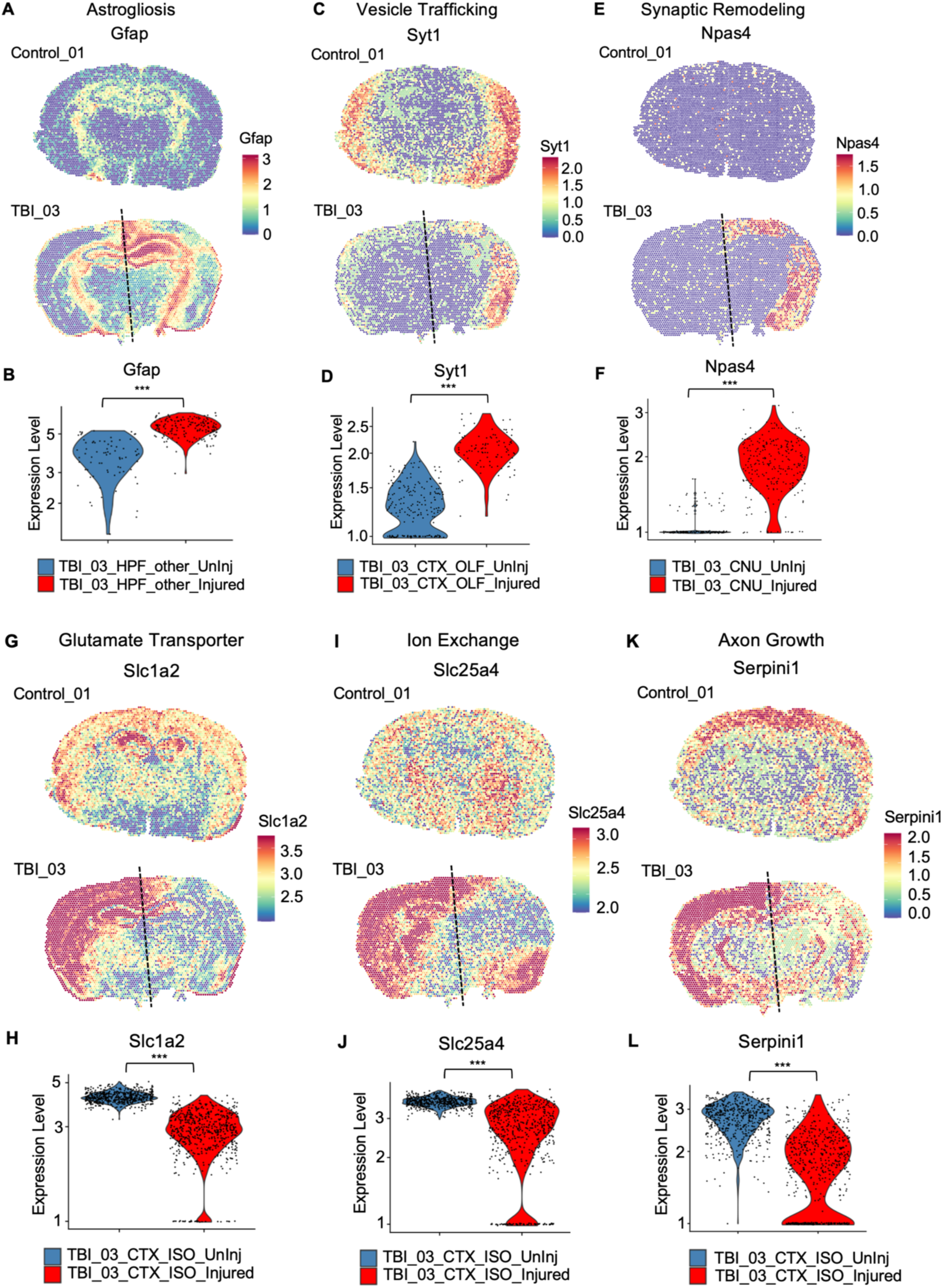
Spatial distribution of key functional marker genes altered post TBI. **A, C, E, G, I, K** Spatial mapping of selected marker genes representing six key functions affected by TBI in Control_01 and TBI_03 samples. Dashed lines separate the uninjured and injured side of the TBI_03 brain. The marker genes and their associated functions are: *Gfap* (astrogliosis), *Syt1* (vesicle trafficking), *Npas4* (synaptic remodeling), *Slc1a3* (glutamate transport), *Slc25a4* (ion exchange), and *Serpini1* (axon growth). Control_01 and TBI_03 samples were integrated using CCA, and for each gene, the gene expression levels are shown using a consistent scale bar across two samples. **B, D, F, H, J, L** Six violin plots illustrating the differences in expression levels of six marker genes in A, C, E, G, I, K, each comparing a selected brain region between the injured and uninjured sides of the TBI_03 sample.

*Syt1* encodes synaptotagmin-1, a calcium-sensing synaptic vesicle protein that regulates neurotransmitter exocytosis; it is critical for synaptic vesicle trafficking. Previous studies have reported that *Syt1* is significantly downregulated following TBI, which could serve as an adaptive response to reduce calcium sensitivity and protect against excitotoxicity following TBI [33]. Our spatial mapping revealed high *Syt1* expression in the cerebral cortex (except the top region of isocortex and hippocampus region) and cerebral nuclei of the control; compared to the control, there was a marked reduction in the same region of TBI samples, especially the uninjured side and the isocortex of the injured side (Fig. 2C). Among the cortical regions, olfactory cortex showed the most pronounced difference in *Syt1* expression between the injured and uninjured sides (Fig. 2D).

*Npas4* is an activity-dependent transcription factor that is rapidly induced by synaptic stimulation. It plays a critical role in synaptic remodeling by activating genes that promote the formation of inhibitory synapses, maintaining the balance between excitation and inhibition in neural circuits [34]. As a marker for synaptic remodeling, *Npas4* was expressed at uniformly low levels across the brain of the control but showed markedly elevated expression in the cortex region on the injured side of TBI samples (Fig. 2E). Notably, whereas the injured hemisphere’s cerebral cortex (except the hippocampus region) consistently displayed increased *Npas4* expression, levels were low at the injury site itself at the top right hemisphere of the TBI_03 sample. There was a significant increase in *Npas4* expression distribution between the cerebral nuclei region on the injured compared to uninjured sides of the TBI brain (Fig. 2F).

For the remaining three marker genes, *Slc1a2* (which encodes a glutamate transporter), *Slc25a4* (which encodes an ion exchanger), and *Serpini1* (which encodes an axon growth factor), we observed similar spatial expression patterns across the samples. These three genes were expressed in a symmetric pattern with relatively high gene expression across the control sample: *Slc1a2* is highly expressed at isocortex and hippocampus region; *Slc25a4* is highly enriched at thalamus and hypothalamus; *Serpini1* is highly expressed at top part of the isocortex (Fig. 2G, I, K). Expression of these genes in TBI samples was decreased relative to the control at the injury site on the right top hemisphere (Fig. 2G, I, K).

Neuronal function requires both neurotransmitter expression and ion exchange. *Slc1a2* encodes Glt-1, which is responsible for clearing the neurotransmitter glutamate from the extracellular spaces in the brain and central nervous system [35], and this regulation is known to be disrupted following TBI, leading to secondary injury processes [36]. Our gene expression analysis provides compelling evidence of this dysregulation: On the injured side of the TBI brain, there was clear downregulation of *Slc1a2* expression; almost the whole injured side of the TBI_03 sample had lower levels of this mRNA than did the control brain (Fig. 2G). When comparing the isocortex regions of both sides of the injured brain, we observed that *Slc1a2* expression was markedly lower on the injured side compared to the uninjured side (Fig. 2H).

*Slc25a4* encodes Ant1, a mitochondrial protein essential for the exchange of ADP and ATP across the mitochondrial membrane [37]. This exchange supports cellular energy supply and contributes to the regulation of ion gradients and electrical properties critical for both mitochondrial function and neuronal signaling. Notably, such regulation of ion exchange can be disrupted after TBI due to impaired energy production and cellular damage [38]. Our spatial mapping of *Slc25a4* gene expression showed a reduction following TBI, which is consistent with this previous finding. The decrease in expression of *Slc25a4* mirrors the pattern observed for *Slc1a2*, most of the injured side shows reduced gene expression, except for the cortical subplate and olfactory cortex regions (Fig. 2I, J).

*Serpini1* encodes the Neuroserpin protein, which is secreted by axons in the brain and plays a critical role in regulating axonal growth and promoting synaptic plasticity [39]. Following TBI, neuroinflammation inhibits axon regeneration by releasing cytokines and toxins that impede neuronal regrowth and by contributing to the formation of a glial scar, which contains molecules that physically and chemically obstruct axon extension. This inflammatory environment impairs the axon growth process [40]. The spatial expression pattern of *Serpini1* is interesting pattern: while Serpini1 expression remains low in the hippocampus on both sides, it is notably high in the cortex of the uninjured side but markedly reduced in the cortex following injury. (Fig. 2K, L).

The evaluation of the expression patterns of the six markers of key cellular processes provided a valuable baseline for assessing TBI severity and enabled comparisons between samples. By applying a spatial transcriptomic approach to map known marker genes, we can quantify the extent of disruption by comparing expression levels between control and TBI samples as well as between the injured and uninjured sides of the brain. Differentially expressed gene analysis may reveal latent genes, potentially identifying novel markers for TBI.

### Spatial Relationships between Genes Define Pathways Unique to TBI

Examining the spatial gene expression of single marker genes provides valuable insights into how specific cellular processes are disrupted after TBI, but a more comprehensive approach is necessary to resolve the spatial patterns of biological processes. The spatial relationships of multiple genes have never been investigated in the TBI brain. By focusing solely on individual gene spatial plots, only differences at the level of single genes and single spots can be observed, without accounting for the interdependence among genes or spatial regions [41]. To better understand these relationships, it is important to investigate how genes are co-expressed and how their expression patterns correlate within and across brain regions. We developed a workflow that quantitatively measures gene-gene relationships and identifies co-expressed gene modules. This approach enables us to visualize the spatial distribution of these modules and quantitatively assess their differences between the injured and uninjured sides of the brain. The enrichment scores for those modules could also serve as an injury score to quantitatively measure the severity of TBI.

The co-expressed gene modules, each representing a set of spatially variable genes with a common set of spatial patterns, were identified using the Giotto package [41]. In brief, we first identified the spatially variable genes and then compared their expression patterns across neighboring cells or tissue regions. By smoothing the data to account for spatial context, genes with similar expression trends were grouped together into modules [41]. We applied this approach to one of the TBI injury samples (TBI_03) and identified 14 distinct gene modules (Fig. 3A-I, Additional file 1: Fig. S3A-E). The number of genes in each module varies from 10 to 100 genes (a detailed list of genes in each module is provided in Additional File 1: Table 1). A heatmap of pairwise spatial correlations of gene expression revealed that genes within each module are strongly co-expressed in space (Additional file 1: Fig. S3G), suggesting coordinated spatial regulation. We then examined the genes within each module and assigned names to the modules based on the identities and functional roles of their constituent genes. The module scores, calculated as the average expression of all genes within each module in each spot on the spatial map, were spatially plotted. Among these, the *Neuronal Activity and Synaptic Plasticity* module, the *Stress and Immune Response* module, and the *Gliosis* module were expressed differently on the injured side than on the uninjured side of the sample (Fig. 3A-C), suggesting that these modules could be injury-specific.

**Figure 3.**
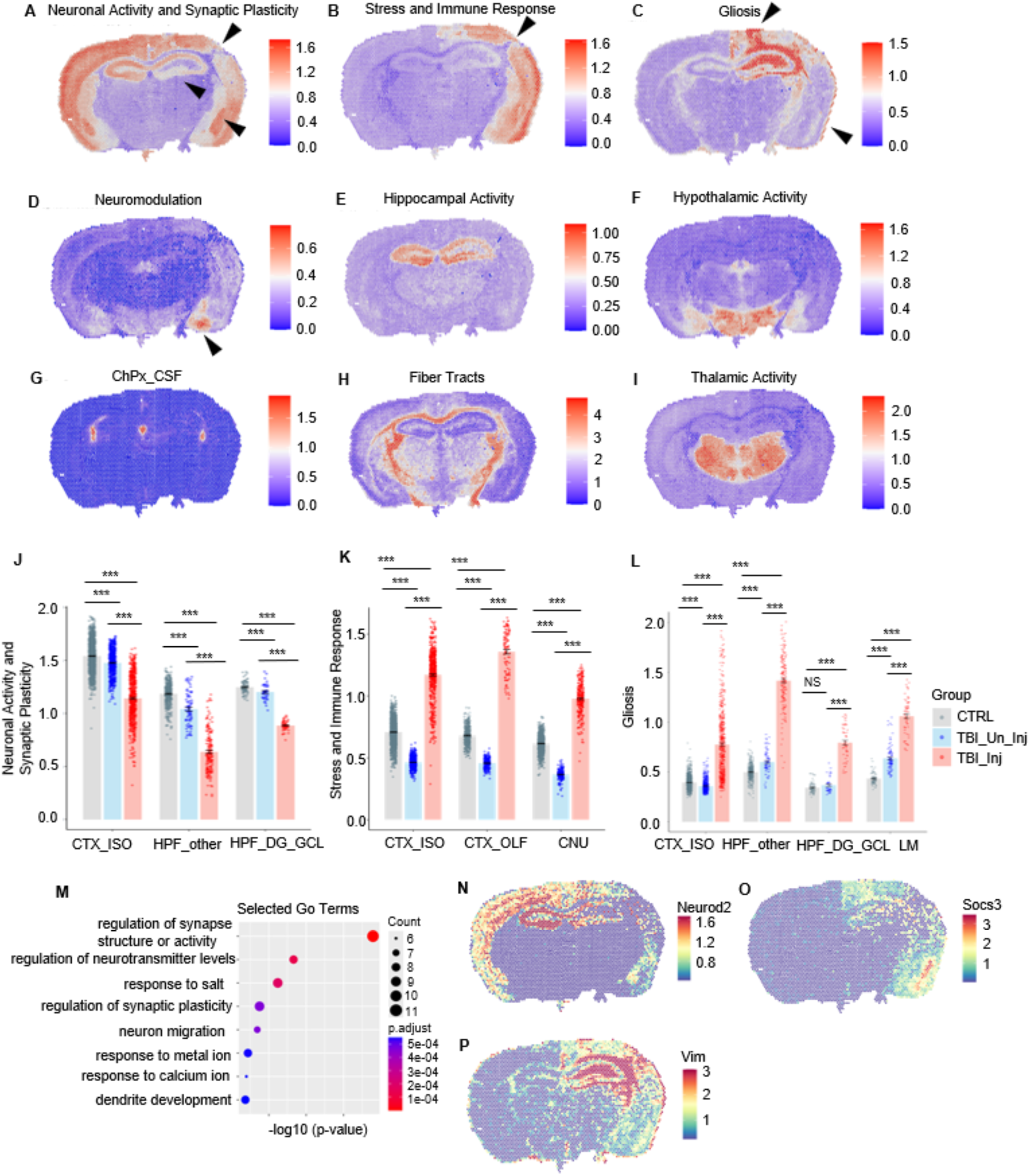
Gene modules highlight key altered biological functions post TBI. **A -I** Spatial distribution of enrichment scores for key gene modules (metagene clusters) across TBI_03 brain tissue sections. A total of 14 gene modules were identified using Giotto algorithm [41]. The nine most representative modules are shown, with each spot indicating the log-normalized average expression of all genes in the module; individual scale bars are used for each module. See Supplementary Fig. 2E-H for additional modules. **J - L** Bar plots of the distribution of module scores for **J**) the *Neuronal Activity and Synaptic Plasticity*, **K**) *Stress and Immune Response*, and **L**) *Gliosis* modules at each spot for selected paired regions on the injured (red) and uninjured (blue) sides of the TBI_03 brain, as well as the region from the Control_01 sample. Each plot includes three or four regions, with each dot representing a single spot from the indicated region. Wilcoxon rank-sum test p-values for each comparison are shown on the plots, *** indicates p < 0.001, while NS denotes a non-significant difference (p ≥ 0.05). Detailed statistical results are provided in Additional File 1: Table 4. **M** Dot plot of manually selected top eight GO biological process terms significantly enriched in the gene set of the *Neuronal Activity and Synaptic Plasticity* module. **N - P** Spatial mapping of gene expression for representative genes from key modules: **N**) *Neurod2* (*Neuronal Activity and Synaptic Plasticity* module), **O**) *Socs3* (*Stress and Immune Response* module), and **P**) *Vim* (*Gliosis* module).

The *Neuronal Activity and Synaptic Plasticity* module is highly expressed in the isocortex, cortical subplate, and hippocampus regions on the uninjured side of the TBI_03 brain (Fig. 3A). In contrast, the module score is relatively low in the isocortex on the injured side, particularly at the injury site in the somatosensory cortex (top right hemisphere, highlighted with black arrow) (Fig. 3A). Similarly, enrichment score is reduced in the hippocampus of the injured side, especially within the dentate gyrus granule cell layer, while remaining high in the cortical subplate (amygdala, bottom right of the cortex, highlighted with black triangle) (Fig. 3 A, J). The *Stress and Immune Response* module is highly expressed in the cortex and cerebral nuclei on the injured side, particularly in the superficial layer of the cortex (Fig. 3B). However, module scores are low at the somatosensory isocortex which is pinpoint with the black triangle. In contrast, this module is expressed at low levels on the uninjured side of the brain (Fig. 3K). Notably, the *Gliosis* module is predominantly localized to the hippocampus, excluding the cornu Ammonis pyramidal layer and the dentate gyrus granule cell layer, and in the leptomeninges and a portion of the isocortex situated above the hippocampus on the injured side (Fig. 3C). The module scores on the injured side are consistently higher than on the uninjured side (Fig. 3L). We also observed an asymmetric pattern in *Neuromodulation* module comparing the injured and uninjured hemisphere. Significant higher module scores were observed in the right amygdala (Fig. 3D). The left and right amygdala have distinct functional roles: the left amygdala is more involved in modulating emotional responses and maintaining long-term emotional regulation, whereas the right amygdala is primarily associated with rapid emotional reactions, such as acute fear and stress reactivity [42, 43]. Therefore, it is reasonable that neuromodulation gene expression is elevated in the right (injured) amygdala, as TBI is likely to trigger acute emotional and stress responses mediated by this region, resulting in hemisphere-specific changes in gene regulation.

We further computed and spatially plotted module scores for the same gene sets identified in the TBI_03 gene modules across one control sample and two additional TBI samples (Additional file 1: Figs. S5, S6). In the control sample (Control_01), module score plots generally exhibited a symmetric pattern, as expected (Fig. S5, S6). However, the *Neuromodulation* module in Control_01 displayed slight asymmetry, with elevated scores observed on the uninjured side near the amygdala (Additional file 1: Figs. S5B). In contrast, the TBI samples (TBI_01 and TBI_02) showed *Neuromodulation* module patterns similar to those of TBI_03, with high expression localized to the amygdala on the injured side (Fig. 3D; Additional file 1: Figs. S5B). This observation aligns with hemispheric lateralization of the amygdala mentioned earlier: high neuromodulation gene expression in the left (uninjured) amygdala is associated with long-term emotional regulation in controls, whereas elevated expression in the right (injured) amygdala of TBI_02 and TBI_03 likely reflects acute fear and stress reactivity resulting from injury [42]. For the *Stress and Immune Response* module, both TBI samples exhibited the expected elevated expression in the cortex and cerebral nuclei on the injured side, similar to observations in the TBI_03 module (Fig. 3B; Additional file 1: Figs. S4B, E, H). The *Gliosis* module in TBI_02 mirrored the pattern seen in TBI_03, with high module scores localized to parts of the leptomeninges and hippocampus, excluding the cornu Ammonis pyramidal layer and dentate gyrus granule cell layer, on the injured side, although expression was not as pronounced as in TBI_03 (Figs. 3C, Additional file 1: Figs. S4C, F, I). In contrast, the *Gliosis* module in TBI_01 did not match the TBI_03 pattern. For TBI_01, the most notable difference in module expression was observed in the fiber tracts on the injured side (Additional file 1: Figs. S4C, F). Since TBI_01 was collected three hours post-injury, whereas TBI_02 and TBI_03 were collected at 24 hours post-TBI, this difference is likely due to the temporal dynamics of astrocyte activation, which typically occurs within a day following injury [44]. Thus, it is reasonable to observe reduced astrocyte activation gene expression in the TBI_01 sample. Additionally, for the *Neuronal Activity and Synaptic Plasticity* module, both TBI_01 and TBI_02 showed very symmetric patterns, with only a slight reduction on the injured side (Additional file 1: Figs. S4A, D, G). This suggests that the effects on neuronal activity and synaptic plasticity are largely confined to the immediate injury site.

By identifying co-expressed gene modules specific to injury, key biological processes that are altered following TBI were identified. The genes within each module show correlated expression and are co-localized spatially, providing a comprehensive view of how groups of genes work together in response to injury. By spatially mapping the module scores, we were able to pinpoint the exact anatomical regions in the TBI brain where these changes occur. Differences between injured and uninjured regions can be quantitatively assessed and visualized.

To validate our module identities, we performed Gene Ontology (GO) term analysis on each gene set [45]. For the *Neuronal activity and Synaptic plasticity* module, significant terms included regulation of synapse structure or activity, regulation of neurotransmitter levels, regulation of synaptic plasticity, and neuron migration (Fig. 3M), all of which are functionally relevant to the module’s biological annotation. It is worth noting that the *Gliosis* module was enriched for several GO terms that are related to the defense response to virus: response to virus, negative regulation of viral process and response to type II interferon (Additional file 1: Fig. 3F). This aligns with the previous finding that the neuroinflammatory response in mice after TBI also triggers the transcriptional activation of endogenous retroviruses [46]. Investigating the relationship between the response to retroviruses and to TBI represents a promising direction for future research.

We evaluated select individual genes from the highlighted modules. *Neurod2*, which is associated with neuron development, showed the expected expression: Low on the injury side compared with the control side (Fig. 3N) [47]. *Socs3*, one of the gene from *Stress and Immune Response* module, is an important negative regulator of cytokine signaling and is crucial in regulating immune response and inflammation [48]. Its gene expression is high at the isocortex, striatum and olfactory cortex, align with the distribution of the *Stress and Immune Response* module score (Fig. 3B, O). *Vim*, like *Gfap*, encodes an intermediate filament protein strongly upregulated in reactive astrocytes after TBI [49]. Its expression is elevated in the hippocampus, except in the cornu Ammonis pyramidal layer, and in the leptomeninges (Fig. 3P), mirroring the spatial pattern of the *Gliosis* module (Fig. 3C). These observed gene expression patterns are consistent with the corresponding gene module plots, further confirming the accuracy of our spatial analyses.

### High-Resolution Spatial Epigenetic Profiling Reveals Fine Regulatory Architecture

After investigating transcriptomic changes following TBI, we sought to understand the potential regulatory mechanisms underlying these differences in gene expression. To address this, we incorporated spatial epigenetic data into OmicGlaze. Our spatial ATAC-seq workflow enables the *in situ* capture of open chromatin regions while preserving spatial information, thereby providing biological insights into how local regulatory landscapes shape region-specific gene expression (Fig. 4A) [3]. First, Tn5 transposase is pre-loaded with specific adapters to allow integration into open chromatin regions where DNA is not tightly packed by histones. These accessible regions often correspond to active regulatory elements such as promoters and enhancers. After transposition, spatial barcoding is performed using microfluidic devices that deliver unique DNA barcodes into the tissue, which are incorporated into accessible chromatin fragments. This preserves the spatial information of the captured fragments.

**Figure 4.**
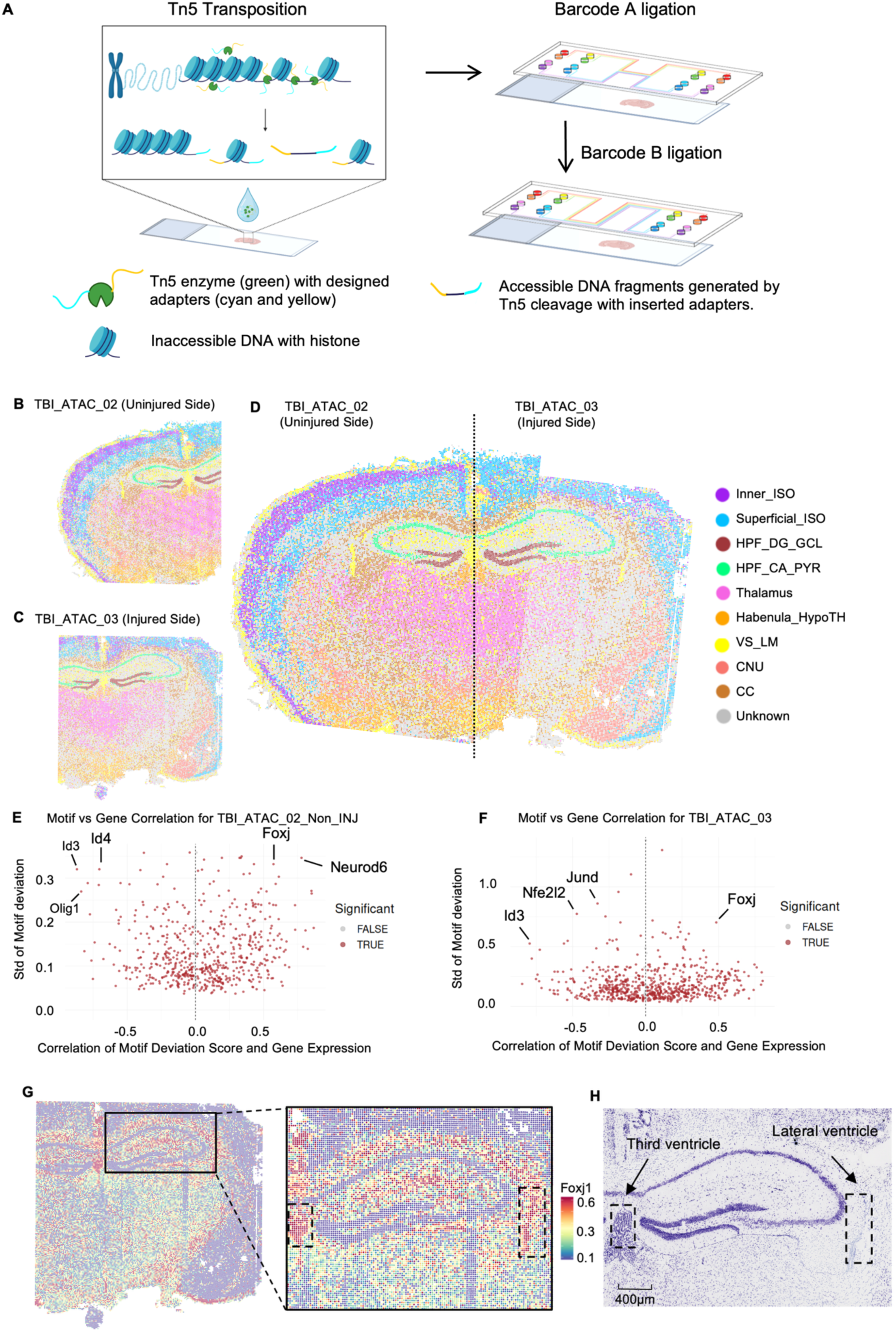
The spatial ATAC-seq workflow profiled chromatin accessibility at a high resolution of 15 μm. **A** Schematic of the spatial ATAC-seq workflow. **B** Spatial distribution of clusters in ATAC-seq data mapped to the uninjured side of the TBI brain (TBI_ ATAC_02_Non_INJ). **C** Spatial distribution of clusters in ATAC-seq data mapped to the injured side of the TBI brain (TBI_ ATAC_03). **D** Integrated view of the uninjured and injured sides for comprehensive comparison. Significant clusters are highlighted with abbreviations: Inner_ISO, inner layer of isocortex; Superfical_ISO, superficial layer of isocortex; HPF_DG_GCL, hippocampal formation dentate gyrus granule cell layer; HPF_CA_PYR, hippocampal formation cornu ammonis pyramidal layer; Habebula_HypoTH: habenula and hypothalamus; VS_LM, ventricular system and leptomeninges; CNU, cerebral nuclei; CC, corpus callosum. **E** Volcano plot showing the correlation between motif scores and gene scores for transcription factors in the TBI_ATAC_02_Non_INJ sample. The y-axis displays the standard deviation of the motif score for each individual motif across spots. Each dot corresponds to a gene; genes with p-values > 0.05 are labeled in red, and those with p-values ≤ 0.05 are labeled in grey. **F** Volcano plot showing the correlation between motif scores and gene scores for transcription factors in the TBI_ATAC_03 sample. **G** Spatial mapping of motif deviation scores for the Foxj1 motif. Each dot reflects the degree to which chromatin accessibility at Foxj1 motif sites at a given spot deviates from the expected background; higher values indicate greater accessibility. A zoomed-in view of the hippocampus on the right highlights the high spatial resolution of the technique. Dashed box on the left indicates the third ventricle, box on the right indicates the lateral ventricle. **H** Nissl-stained image of the adjacent slide from the TBI_ATAC_03 sample, with a zoomed-in view of the hippocampal region corresponding to panel H. The dashed box on the left indicates the third ventricle, and the box on the right indicates the lateral ventricle. Scale bar represents 400 µm.

The four tissues analyzed using spatial ATAC-seq came from the same brains as those used in the spatial transcriptomics experiments: Control_ATAC_01, TBI_ATAC_01, TBI_ATAC_02_Non_INJ and TBI_ATAC_03. For TBI_ATAC_02_Non_INJ and TBI_ ATAC_03, sections were taken from adjacent slides, with ATAC_TBI_02_Non_INJ corresponding to the uninjured side and ATAC_TBI_03 to the injured side. Each mapped a 5-mm region of the tissue at a high spatial resolution of 15 μm per pixel. This high-resolution spatial mapping enabled us to examine chromatin accessibility patterns across different brain regions and injury states at cellular level. We examined the number of fragments, fragments sizes and transcription start site enrichment score as quality control metrics, and they all showed that our sample were of good quality (Additional file 1: Fig. 7, Table 3).

We next performed unsupervised clustering on each of the four samples. Several clusters that clearly correspond to anatomical regions of the mouse brain were observed in all four samples, as identified based on transcriptomic profiles: superficial isocortex, inner isocortex, thalamus, cornu Ammonis pyramidal layer and dentate gyrus granule cell layer of hippocampus formation region, and cerebral nuclei (Fig. 4B and C, Additional file 1: Fig. S8A-F). On the uninjured side (TBI_ATAC_02_Non_INJ), the two layers of isocortex are clearly distinguished by their respective clusters, whereas on the injured side (TBI_ ATAC_03), the inner isocortex layer appears disrupted, with a mixing of superficial and inner isocortex clusters. Given that these samples are derived from adjacent sections, the two isocortex clusters were manually combined to achieve an integrated, high-resolution map of transcriptomic features across the entire mouse brain (Fig. 4D). This approach offers a comprehensive spatial overview and enhances our ability to resolve the epigenetic change following TBI at fine anatomical scales.

We next performed peak calling and transcription factor motif deviation enrichment analysis using ChromVAR [50]. The transcription factor motif scores quantify how much the chromatin accessibility of motif-containing regions in a cell (or in our case, a spot on the microfluidic device) deviates from what would be expected by chance. Since this deviation score is calculated on a per-cell basis, in our spatial context, we were able to plot the motif enrichment spatially. The correlations between gene scores and motif deviation scores were determined for the TBI_ATAC_02_Non_INJ and TBI_ATAC_03 pair (Fig. 4E, F). Each point represents a transcription factor motif, with the *x*-axis showing the Pearson correlation between its motif deviation score and the gene score of the corresponding gene, and the *y*-axis indicating the standard deviation of motif deviation across spots. A high positive correlation indicates that both the gene and the corresponding transcription factor binding site are highly accessible in the analyzed sample, suggesting that the transcription factor is an activator of expression of itself under the evaluated condition. Conversely, a low gene score and a low motif deviation score suggest that the transcription factor acts as a repressor of itself. In the meantime, a high standard deviation of motif deviation scores means that the motif accessibility varies substantially across spots in the sample, potentially representing transcription factors with dynamic or region–specific regulatory activity. Some of the transcription factors in the TBI_ATAC_03 sample show greater than 0.5 standard deviation of motif deviation scores, while none of the transcription factors reaches a standard deviation higher than 0.5. This shows that on the injury side, an increased variability and stronger motif–gene correlations post-TBI presents, meaning gene regulatory network is being reprogrammed at an injury-specific manner.

From this analysis, we identified several transcription factors that had either strong positive or strong negative correlations with their corresponding gene and high standard deviation for their motif deviation score in the TBI_ATAC_03 sample. For example,׈*Foxj1*, which is known as a master regulator of motile ciliogenesis and is directly required for the differentiation of ciliated cells by activating genes essential for cilia formation in mice [51], had strong positive correlation between its gene score and motif deviation score, consistent with its role as an activator. In contrast, Id3, which functions as a dominant-negative repressor of other basic helix-loop-helix transcription factors to inhibit cellular differentiation [52]. It had strong negative correlation between its motif deviation and gene score in both TBI samples, supporting its function as a repressor. We spatially plotted the motif deviation score for *Foxj1* to demonstrate the high-resolution mapping capability of our approach (Fig. 4G). The plot revealed that deviation scores are high in the ventricular system which are ciliated cells located in the brain consistent with its activator role of motile ciliogenesis. The zoomed-in view also highlights the fine spatial resolution that clearly distinguishes the cornu Ammonis pyramidal layer and the dentate gyrus granule cell layer of the hippocampus, both of which display relatively low motif deviation scores compared to surrounding regions. The two black rectangles also emphasize the increase motif deviation score at the third ventricle and the lateral ventricle with fine resolution. These regions were shown for reference in the adjacent Nissl-stained image (Fig. 4H). This epigenetic workflow thus offers a detailed view of chromatin accessibility in the brain and provides a powerful tool for studying epigenetic changes that regulate gene expression following TBI.

### Spatial Dissection of Gene Regulatory Mechanisms in the TBI Brain

Using high-resolution spatial epigenetic profiling and the annotation of mouse brain regions, we identified factors that may regulate changes in gene expression after TBI within the tissue context. We first calculated the motif deviation scores by jointly analyzing the ATAC-seq data for samples from the uninjured and the injured side of the same TBI brain (TBI_ATAC_02_Non_INJ and TBI ATAC_03), resulting in motif scores calculated based on the overall background. We then identified differentially active motifs across the two samples and plotted with a volcano plot (Additional file 1: Fig.S8G). Notably, we highlighted Atf3, a transcription factor known to modulate stress and immune responses [53]. It showed a high positive log2 Fold change meaning its motif accessibility was markedly increased in the injured side of the TBI sample relative to the uninjured side, suggesting its potential regulatory involvement in injury-induced chromatin remodeling. We visualized the spatial distribution of the Atf3 motif deviation scores across both samples and confirm that, in the injured side of the TBI brain (TBI_ATAC_03), the isocortex, olfactory cortex, cortical subplate, and cerebral nuclei had markedly higher Atf3 motif deviation scores than the corresponding regions in the uninjured side (TBI_ ATAC_02_Non_INJ) (Fig. 5A). This pattern is highlighted in the zoomed-in views of the hippocampus and parts of the isocortex (Fig. 5B and C).

**Figure 5.**
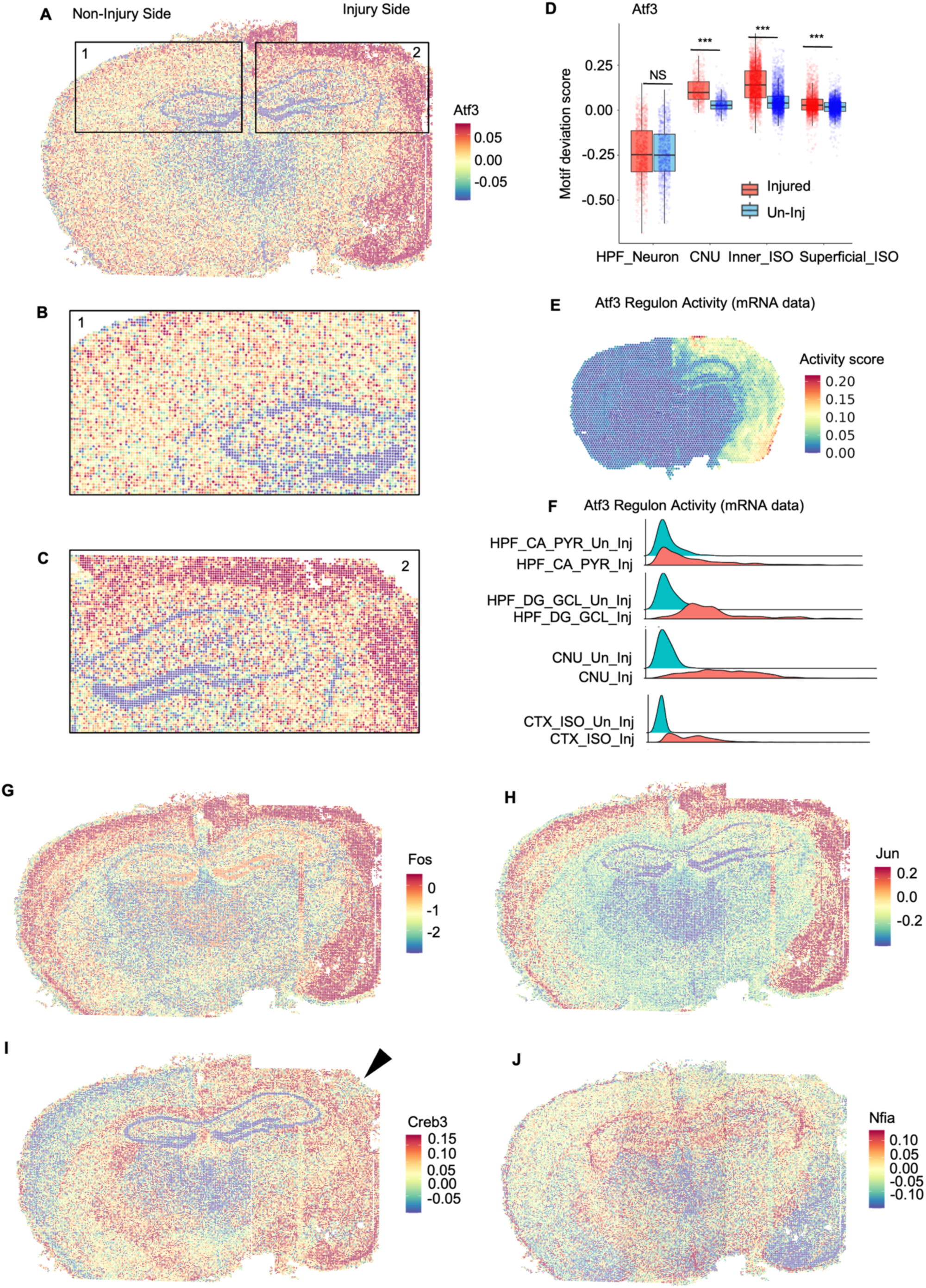
Epigenetic analysis of transcription factor–driven gene regulation. **A** Spatial motif deviation scores for the Atf3 motif, shown for both the uninjured (TBI_ ATAC_02_Non_INJ) and injured (TBI_ ATAC_03) sides of the brain. Motif deviation scores were calculated using integrated data of all samples, and a shared color scale bar was used for comparison. Red spots indicated that the Atf3 motif was located within an open chromatin region. Blue spots showed the inaccessible motifs which recognized by Atf3 transcription factor. **B** Enlarged view of the Atf3 spatial motif deviation scores at the uninjured side (TBI_ ATAC_02_Non_INJ), highlighting the contralateral hippocampus and isocortex regions. **C** Enlarged view of the Atf3 spatial motif deviation scores at the injured side (TBI_ ATAC_03), highlighting the hippocampus and part of the isocortex regions. **D** Boxplots comparing Atf3 motif deviation score distributions between paired regions in the injured and uninjured hemispheres. Four region pairs were analyzed: HPF_Neuron, CTX_SP, Inner_ISO, and Superficial_ISO. The HPF_Neuron pair shows no significant difference (t-test p = 5.2 × 10⁻¹; NS), whereas the other three pairs exhibit highly significant differences (p < 2.2 × 10⁻¹⁶; ***, representing the minimum calculable value in R). **E** Spatial plot of the Atf3 gene regulatory network activity, inferred from spatial mRNA-seq data. Color scale represents the combined expression of all the downstream genes in the Atf3 family. **F** Atf3 regulon activity scores between paired regions on the uninjured and injured sides of the TBI_03 brain. Blue denoted regions on the uninjured side, and red denoted regions on the injured side. **G - J** Spatial motif deviation scores for other transcription factors G) Fos, H) Jun, I) Creb3, and J) Nfia.

To quantify differences in motif deviation scores across paired regions from the injured and uninjured sides, we compared the distribution of motif deviation scores for each spot in each region using a boxplot (Fig. 5D). For the cornu Ammonis pyramidal layer and dentate gyrus granule cell layer of hippocampus formation region, no significant difference was observed, whereas cerebral nuclei, superficial, and inner isocortex regions had higher motif deviation scores in the injured brain than the uninjured brain. Next, we applied the SCENIC package to identify the Atf3 regulon, which includes genes co-expressed with Atf3 and carrying Atf3 transcription factor motifs in their regulatory regions [54]. Genes identified in the regulon is shown in Additional file 1: Table 5. This gene set includes well-known immune-related genes such as *Cd14, Ifit3*, and *Ccl3* [55–57], as well as classic stress-response genes including *Egr1, Ddit3*, and *Bdnf* [58–60]. We spatially plotted out the Atf3 regulon activity, which measures the combined expression of genes in the regulon. The overall expression for those targeted genes is high in the isocortex, striatum and olfactory cortex which highly overlap with the increased Atf3 motif opening in these regions (Fig. 5A, E, F). The overlap in spatial patterns indicates that Atf3 functions as a central regulator coordinating both immune activation and stress-response programs post TBI.

Since Atf3 is a member of the Activator Protein-1 (AP-1) family, we additionally examined two other members of this family, Fos and Jun, which regulate expression of genes involved in cell survival and inflammation [61]. Both of these transcription factors have similar spatial motif deviation patterns: The isocortex, striatum and olfactory cortex showed high motif activity on the injured side, whereas on the uninjured side those regions had lower motif deviation scores compared to the injured side (Fig. 5G. H).

Our method also detected other spatially variable transcription factors, not related to AP-1 family. For instance we identified Creb3, a transcription factor activated by endoplasmic reticulum stress that regulates adaptive and apoptotic gene expression [62]. In the uninjured side, both layers of the isocortex had low Creb3 motif deviation scores, but on the injured side, higher motif deviation was observed in the isocortex, olfactory cortex, cerebral nuclei, and cortical subplate (Fig. 5I). These are regions likely to experience injury-related stress. Interestingly, at the somatosensory cortex (point by the black arrow in the figure), the deviation score was lower than in surrounding regions (Fig. 5I).

Finally, we also identified transcription factors that were not changed post TBI. Nfia, a transcription factor involved in regulation of myelination [63], did not differ between control and TBI samples in spatial motif deviation plot (Fig. 5J). For validation, we look at the gene expression for one of its targeted gene, *Mbp* which is a key gene involved in myelin sheath formation [64]. Our spatial analysis of *Mbp* revealed no significant differences in expression between samples, align well with the motif deviation score pattern for Nfia (Additional file 1: Fig.S8H). This consistence between the transcription factors motif accessibility and its targeted gene expression validates our approach as a robust method for benchmarking transcription factor regulation after TBI.

Using a spatial multi-omics approach, OmicGlaze revealed region-specific alterations in both transcriptomes and epigenomes following TBI. The spatial transcriptomic patterns of the genes offered valuable insights for the local biological pathways. We found that neuronal activity and synaptic plasticity, stress and immune response, and gliosis were activated during the acute phase of injury. For the first time, we mapped the motif activity of Atf3, a master transcription regulator, in regulating the stress response post TBI. Differences in motif deviation scores between the injured and uninjured hemispheres explain why genes in the isocortex are more responsive to injury, as their epigenomes are remodeled by Atf3 and its AP-1 family partners Jun and Fos.

## Discussion

In this study, we developed *OmicGlaze*, a spatial toolbox to elucidate the pathogenesis of TBI-induced secondary injuries by defining transcriptomic and epigenomic changes in layer-specific subregions. Our method quantitatively surveyed entire coronal sections (TBI versus Control, Peri-injury versus injury-center plane, Contralateral versus Ipsilateral), revealing molecular hallmarks for the initiation and propagation of TBI-induced secondary injuries that cannot be detectable by conventional histopathology. *OmicGlaze* is a new computational workflow to explore both spot-spot differences and also gene-gene relationships. It allows numeric comparison between different spatial datasets by gene module scores (Figure 3). The calculated scores of these gene modules offer a quantitative measure of TBI severity, as a new standard tool in investigating spatiotemporal changes in TBI.

Our approach enables a comprehensive, multi-omics analysis that directly links region-specific changes in gene expression to underlying chromatin accessibility and transcription factor activity. Our experimental workflow incorporates several optimized procedures tailored specifically for TBI study. Fresh frozen, unfixed mouse brain tissues were processed using optimized flash-freezing protocols to preserve molecular integrity, informed selection of anatomical planes exhibiting significant TBI-induced alterations, and a sectioning strategy to ensure comparable levels across samples. For spatial transcriptomics, we used Visium V2 CytAssist platform (11 mm x 11 mm) to capture the entire coronal section of post TBI mouse brains, allowing region-specific analysis in contralateral hemispheres and also peri-injury planes. We established a quantitative scoring metric to evaluate altered biological pathways in spatial data, and offered it as a normalized value to compare between spatial datasets.

We present the first spatially resolved map of gene regulation in traumatic brain injury. Our spatial ATAC-seq experiment was optimized by using the pre-patterned FlowGel system (AtlasXomics), enabling tissue-level coverage (5.5 mm x 5.5 mm) at a high resolution of 15 µm. One single tissue section yielded a total of 48,400 spots, which was 20 times more than our previously spatial ATAC-seq platform [18]. Leveraging this high-quality epigenetic data, we identified region-specific epigenetic activities of the AP-1 transcription factor family that cellular stress and immune response. Within the AP-1 family, we identified Atf3 as a key regulator for stress and immune response post TBI, characterized by elevated chromatin accessibility at its binding motifs in the injured isocortex. Gene regulatory network analysis by SCENIC [54] of the Atf3 regulon revealed increased expression of both Atf3 and 89 target genes in specific regions of the injured hemisphere relative to the uninjured hemisphere. Atf3, along with other transcription factors yet to be identified, represents a high-value therapeutic target. Interventions aimed at modulating these regulators may help restore normal cellular transcriptional states and potentially reverse the gene expression alterations induced by TBI. Together, *OmicGlaze* provides new insights into the acute molecular response to TBI, and demonstrates the power of spatial multi-omics for unraveling the regional and regulatory complexity underlying secondary injury processes.

Future studies will require novel computational integration of spatial ATAC-seq and spatial RNA-seq datasets. Although computational methods to integrate single cell multi-omics data have been established [65], there is a gap in applying such method to spatial data. Experimental technological advances in co-profiling the transcriptome and epigenome with spatial coordinates would be an alternative option [66].

Several limitations of our study were acknowledged. First, the temporal dynamics of TBI were not thoroughly investigated, as our analysis focuses primarily on samples collected at 24 hours post-injury. Future studies incorporating additional samples from strategically selected time points (e.g., 1,2,4,6,24 hours) would enable deep and more systematic insights into the progression of TBI-associated molecular changes over time. Furthermore, although we identified region- and cell type-specific alterations post-TBI, deconvolution of cellular heterogeneity was not performed in this study. Integrating reference single-cell RNA-seq datasets derived from TBI samples would facilitate high-resolution, cell-type deconvolution at the single cell level. The challenge holds in financial limitation in collecting a TBI-specific single cell dataset that at least 20 times larger than the spatial dataset. Existing single-cell references provided at the Allen Institute were derived from healthy mouse brain, therefore lack any disease-cell populations relevant to TBI. As the field progresses, emerging single-cell and spatial technologies will help overcome these limitations and ultimately support new prognostic tools and therapeutic strategies to prevent or mitigate TBI-induced secondary injuries.

## Methods

### Animal model of traumatic brain injury

Two types of TBI models are used in this study. All experimental procedures were performed in accordance with institutional guidelines and approved by the Bloodworks Northwest Research Institute under protocol #111-02 and University of Michigan Institutional Animal Care and Use Committee under protocol #IACUC PRO00012315. For fluid percussion injury (FPI) model, we subjected mice to a FPI using a pendulum device (Custom Design & Fabrication) [67]. C57BL/6J mice (10 weeks old, male) were given 0.5 mg/kg buprenorphine (Wedgewood Connect) before being subjected to a 3-mm diameter craniectomy (approximately 2 mm rostral to Bregma and 2 mm lateral to the sagittal suture) in the right hemisphere under isoflurane anesthesia (2.5-3% flowrate, 0.8 O2 lpm; VetOne). A modified Luer-Lok needle hub (4-mm inner diameter, 20 g (BD) was secured over the exposed, intact dura mater with cyanoacrylate adhesive (Loctite, Rocky Hill). The assembly was secured to the skull with dental cement (Lang Dental). One day after the craniectomy, the injury tube was connected to the fluid percussion cylinder. A toe pinch response was elicited before FPI to ensure that all animals were at the same level of unconsciousness. The mice were subjected to a one-time lateral FPI at 1.9±0.6 atm under isoflurane anesthesia with the head restrained to mimic TBI in most clinical settings. The transient magnitude of the pressure pulse exerted on the dura was measured with a pressure transducer (Custom Design & Fabrication) and oscilloscope (GW Instek). Immediately after TBI, mice were monitored for breathing, and time to righting reflex was recorded. Mice were anesthetized under isoflurane, and their incision was closed with sutures. Mice were injected with sterile PBS via tail vein 30 minutes after injury. At three hours post TBI, mice were euthanized by carbon dioxide inhalation and cervical dislocation.

In the controlled cortical impact (CCI) TBI model, TBI was induced using a CCI device (Impact One™ Stereotaxic CCI Instrument, Leica Biosystems, #39463920) [68]. Adult C57BL/6J MitoTimer/Thy1 Cre mice (8 months old, male) were anesthetized with 2-4% isoflurane in oxygen and secured in a Vernier stereotaxic frame (Leica Biosystems, #39463201). Following a midline scalp incision and reflection of the periosteum, a 4.0-mm diameter craniectomy was performed over the right parietal cortex, lateral to the sagittal suture and between lambda and bregma, with care to preserve dural integrity. The exposed cortical surface was subjected to a single controlled impact using a 3.0-mm diameter flat-tip electromagnetic impactor programmed to deliver 1.0-mm tissue deformation at 3.0 m/s velocity with a 100 ms dwell time. These parameters produce a moderate-severity focal brain injury with reproducible cortical and hippocampal damage. Post-impact, the scalp incision was closed with interrupted sutures, and subcutaneous carprofen (5.0 mg/kg), and saline was administered. Animals recovered from anesthesia in a temperature-controlled recovery environment and were monitored until fully ambulatory before being returned to housing. Sham-operated controls received identical anesthesia, surgical preparation, and craniectomy without cortical impact to control for surgery-related effects. The brains were harvested 24 hours after injury.

### Tissue collection

The fresh mouse brains were soaked in 30% sucrose DPBS solution with RNase inhibitor (New England Biolabs, Inc., #M0314L) added to a final working concentration of 1 unit/µL. The brains were temporarily stored in 15 mL conical tubes on wet ice for up to 4 hours before flash freezing. Excess liquid on the brains was removed using lint-free clean room wipes (Uline, #S-21888). The whole brains were flash frozen by holding the brain cradled in an aluminum foil boat above dry ice in methylbutane vapor for a few seconds until the edges of the tissue turned white, and then submerged in methylbutane for 1–2 minutes to ensure complete freezing. The samples were then placed in zip bags with desiccants. Bags were sealed and stored at −80 °C freezer until sectioning to prevent condensation damage.

### Tissue sectioning

Prior to sectioning, we embedded the tissue in OCT compound, covering regions that included the hippocampus and posterior brain. The anterior portion of the brain was trimmed at the cryostat (Leica, #CM3050S). Sectioning and histology staining were performed at the Molecular Pathology Shared Resource of the Herbert Irving Comprehensive Cancer Center at Columbia University (New York State, USA). Because of natural variations in sectioning orientation, obtaining identical coronal planes across different samples was not feasible. In this study, we primarily used coronal sections taken around the mid-hippocampal region. We placed one coronal brain section, 10-µm thick, on each glass slide (Leica, #3800081), yielding a total of 40 consecutive sections per brain. We stained the slices adjacent to the spatial transcriptomics section with H&E and for Nissl bodies (Figure 1B).

### Spatial mRNA sequencing

We used Visium V2 CytAssist Spatial Gene Expression Reagent Kits (10X Genomics, #1000523) to profile mRNAs from tissue slices. Following manufacturer’s user guide (CG000614 and CG000685), we incubated the tissue slice with the Visium Mouse Transcriptome Probe Set (V1), and then transferred the tissue with probes from standard glass slides to a Visium slide using an CytAssist instrument (Software Version 2.2.0.8). Each Visium slide had a capture area of 11 mm x 11 mm, providing coverage for the entire coronal slice of a mouse brain. The spatial resolution is based on 55-μm diameter spot with a center-to-center distance of 100 μm. The resulting libraries were sequenced on an Illumina NovaSeqX sequencer with pair-end 2x150 bp targeting more than 25,000 reads per spot. Quality metrics, including the total number of reads per sample, fraction reads in spots under tissue, and sequencing saturation, are summarized in Additional file 1: Table 2.

### Alignment and preprocessing of spatial transcriptomics data

We used the Space Ranger software pipeline (10x Genomics, Version 3.1.3) to extract the expression matrix of Visium spatial transcriptomics data. Raw sequencing files (FASTQ format) were aligned to the mouse reference genome (mm10-2020-A) using Space Ranger. We performed manual fiducial alignment using the H&E image in the 10X Loupe software (version 9.0.0, 10x Genomics). We used the “Visium Mouse Transcriptome Probe Set v1.0” in the 10X CytAssist Visium pipeline, following manufacturer’s default protocol. A total of 19,465 gene identities (targeted by 19,763 probes) were present in the final filtered output. The quality control data, including the number of spots under tissue, mean reads per spot, median genes per spot, median UMI counts per spot, and total gene detected, for each Visium chip are summarized in Additional file 1: Table 2.

The expression matrix generated from Space Ranger, including the spatial gene expression matrix and associated spatial metadata were imported into Seurat (version 4.3.0) using Read10X and Read10X_Image functions. We first performed CCA integration on all four samples as described by Seurat (https://satijalab.org/seurat/articles/seurat5_integration) [22]. The data were then normalized using the SCTransform function and a shared nearest neighbor graph was constructed using the top 30 principal components with the FindNeighbors function. Unsupervised clustering was then performed using the Louvain algorithm via the FindClusters. UMAP dimensionality reduction was performed using the top 30 principal components with the RunUMAP function to project the data into a two-dimensional space for visualization (Fig. 1J). The FindMarkers() function was used to identify the marker genes for Control_01 sample for each region, and the heatmap was plotted with the DoHeatmap function. To compare the gene expression between the Control_01 and TBI_03 samples, we first extracted the data for those two samples from the CCA integrated dataset, and we then used the SpatialFeaturePlot function from Seurat to plot the gene expression from the Control_01 and the TBI_03 sample. Violin plots were created using the VlnPlot() function.

### Automated Segmentation and Quantification of Cells Across Brain Hemispheres

Whole-slide H&E images (0.5 µm px⁻¹, Leica Aperio) were partitioned into sixteen non-overlapping tiles (4 × 4 grid) with pyvips-binary (v8.14.2) for BigTIFF input and Pillow (v10.3.0) for JPEG input/output. Tiled images were preserved with EXIF orientation, adjusted with edge-tile dimensions to fit the native field of view, and exported with lossless LZW-compressed BigTIFF and high-quality JPEG tiles. These sixteen tiles produced from one slide constituted the training dataset; tiles were manually annotated for soma/nuclei detection.

Delineation of somata and neurites was carried out with Cellpose-SAM (v4.0.5). A custom model was trained for 100 epochs on the manually annotated 16 representative tiles (learning rate 1 × 10^-4, weight decay 0.1). The trained model was then applied to untiled H&E images to generate uint32 masks for detected soma/nuclei.

Hemispheric boundaries were annotated in ImageJ/Fiji (v1.55) using a custom macro to draw a single vertical partition on each H&E tile. After the line was positioned, the macro generated an 8-bit binary image the same size as the source tile, filled the pixels on one side of the line with foreground (value = 255) and the opposite side with background (value = 0), and saved the result as a TIFF mask.

Generated masks and the corresponding H&E images were imported into CellProfiler (v4.2.8). Hemisphere-specific object counts were obtained with two MaskObjects operations that applied the binary split mask created in ImageJ/Fiji to the Cellpose-identified nuclei, producing left (control) and right (injured) subsets. Objects were retained only when at least 50 % of their pixels overlapped the relevant hemisphere mask; nuclei falling below this threshold were discarded. The numbering of surviving objects was preserved. All object-, image-, and experiment-level measurements were exported through the ExportToSpreadsheet module for downstream analysis.

Quantification tables were parsed in R (v4.5.0) using tidyverse (v2.0.0), ggpubr (v0.6.0), ggstatsplot (v0.10.1) and rstatix (v0.7.2). For each histological Section identifier, counts were summed within hemisphere, yielding one paired observation per Section (n = 5). Paired differences (right − left) were tested with a two-sided Wilcoxon signed-rank test; effect size was expressed as the rank-biserial correlation with a percentile-bootstrap 95 % confidence interval (10, 000 resamples). Visualizations were produced with ggplot2 (v3.5.0) using the “jco” palette.

### Region annotation and analysis

We first utilized CCA to integrate data from two sets of sample pairs: Control_01 with TBI_01 and TBI_02 with TBI_03. Following integration, we applied the FindClusters function with a resolution of 0.5 to perform unsupervised clustering of the regions. Each cluster was then annotated by referencing the Allen Institute Mouse Brain Atlas [23], specifically using Coronal Atlas image 74 to map clusters to their corresponding brain regions. Clusters that did not correspond to any specific region in the atlas were labeled as “Unknown.” To enhance and refine regional annotations, we used Loupe Browser (10X Genomics). We imported the .cloupe files, which were automatically generated from the Space Ranger pipeline, into Loupe. This software allowed us to manually select and annotate spatial spots, which enables us to distinguish between the injured and uninjured sides of the TBI samples. Annotations were exported as CSV files, imported into R, and added as metadata labels the side to the Seurat object. This workflow also enabled us to further refine regional annotations by modifying assignments of spots that did not clearly belong to a specific region.

### Identification of co-expressed gene modules using Giotto

A Giotto object was created directly from the Visium output using the createGiottoVisiumObject function or gene expression information and spatial coordinates were extracted from the Seurat object and utilized as input for the createGiottoObject function [41]. Subsequently, we followed the steps described by the developers of the Giotto package (https://rubd.github.io/Giotto_site/articles/tut10_giotto_spatcoexpression.html). Module scores were added to the TBI_03 Seurat object as metadata, and we plotted the spatial distribution and bar plot distribution using SpatialFeaturePlot function and ggplot function. The enrichGO function in the clusterProfiler package was used for GO enrichment analysis with a qvalueCutoff of 0.05 [45].

### Spatial-ATAC sequencing

Open chromatin was profiled with spatial ATAC-seq technology [18, 69] by AtlasXomics. Each chip had a capture area of 5.5 x 5.5 mm^2^, with a 15 mm spot size, totaling of 48,400 spots. The center-to-center distance of two spots was 25 μm. Following the AtlasXomics protocol (AXO-0455(05)), the tissue was thawed at 37 °C for 5 minutes. Then, it was fixed with 0.2% formaldehyde for 5 minutes and quenched with 1.25 M glycine for 5 minutes. The tissue slice was washed twice with 1 ml of DPBS and then deionized water. The tissue slice was then permeabilized for 15 mins, washed with wash buffer for 5 minutes, and then tagmentation was performed at 37 °C for 30 minutes. After addition of EDTA, the tissue was fixed with 4% paraformaldehyde to prepare for spatial barcoding. For spatial barcoding, the AtlasXomics protocol AXO-0457(08) was used. The tissue was stamped with FlowGel chips containing spatial barcodes using the AtlasXpress clamp for 35 minutes at 37 °C in a humidity chamber. After the gel was removed by incubation at 37 °C in PBS, the ligation solution, containing T4 ligase (lig B) and ligation buffer (lig A), was pooled onto the tissue for 15 minutes to ligate the spatial barcodes onto the tissue after hybridization during the stamping step. Tissue was incubated in NEB buffer and then dipped in H_2_O and dried. These steps were repeated for the second stamp, and the region was lysed with lysis buffer overnight in the AtlasXpress clamp. The lysate was collected and then purified with Zymo DNA Clean & Concentrator-5. The library was amplified according to AtlasXomics Protocol (AXO-0459(05)). The number of cycles performed was determined using the cycle with one-third the intensity of the maximum signal in a qPCR. The library was evaluated using Qubit 4 (Thermo Fisher) and an Agilent D5000 chip on a Tapestation (Agilent, #4150) and sequenced on an Illumina NovaseqX+ with 150 base pair paired-end reads.

### ATAC data analysis

FASTQ files were aligned to the mouse reference genome using Chromap.38, creating BED-like fragment files for downstream analysis. Additionally, microscopy images of the regions interrogated were processed through the AtlasXBrowser software [70] to remove any pixels with no tissue as well as provide metadata for the individual barcode locations.

Spatial ATAC-seq analysis was performed using the ArchR package (version 1.0.2) in R (version 4.3.3). The processed fragment files were converted into Arrow files using the createArrowFiles() function, with a tile matrix added to the object using a genome binning size of 5 kb. Arrow files were converted into a single ArchRProject for downstream analysis. The fragment file was read into ArchR as a tile matrix with a genome binning size of 5 kb, and pixels that not present on the tissue were removed based on the metadata file generated in the previous step. Cell metadata were extracted to remove the pixels that were not on the tissue, based on matching spatial barcodes from the tissue sections with cell barcodes in the ArchR project.

Dimensionality reduction was performed using an iterative Latent Semantic Indexing (LSI) approach (addIterativeLSI(), selecting the top 25,000 variable features and the first 30 LSI dimensions. Clustering was performed using the addClusters() function with a resolution of 0.5; the addUMAP() function was used for visualization in a low-dimensional space. To enhance downstream analyses by smoothing the sparse chromatin accessibility data, imputation weights were calculated with the addImputeWeights() function. The Gene Score model in ArchR was used to calculate gene accessibility scores, resulting in the creation of a gene score matrix for subsequent analyses.

Fragment files were processed using the SnapATAC2 package to perform clustering based on a tile matrix. Fragments were imported using the import_fragments() function, and a tile matrix was generated with a bin size of 5 kb using add_tile_matrix(). The top 25,000 features were selected for downstream analysis. Dimensionality reduction was performed using spectral embedding, and the first dimension, which is often associated with sequencing depth, was excluded. A k-nearest neighbor (KNN) graph was constructed, followed by clustering using the Leiden algorithm with a resolution parameter of 3. Uniform Manifold Approximation and Projection (UMAP) was used to visualize cells in a low-dimensional space. Each cluster was then annotated by referencing the Allen Institute Mouse Brain Atlas [23], specifically using Coronal Atlas image 74 to map clusters to their corresponding brain regions. Clusters that did not correspond to any specific region in the atlas were labeled as “Unknown.”

The gene score data and the clustering information from SnapATAC2 pipeline [71] were loaded back to Seurat v 4.3.0 to map the data back to the tissue section and obtain the spatial visualization. Pseudobulk group coverages were generated according to cluster identities with the addGroupCoverages function, and these were utilized for peak calling with MACS2 via the addReproduciblePeakSet function in ArchR. After creating the background peak set using addBgdPeaks, we utilized chromVAR to calculate per-cell motif activity, referred to as the motif deviation score [50]. This score enabled spatial mapping of motif activity. We ran chromVAR using the addDeviationsMatrix function with the CIS-BP motif set. The resulting motif deviation scores were then imported back into Seurat for visualization.

To assess the relationships between gene scores and motif deviation scores, we used ggplot2 to generate a volcano plot illustrating their correlation. Region specific marker motifs were identified using the getMarkerFeatures (bias = c(TSSEnrichment, log10(nFrags), testMethod = Wilcoxon) and getMarkers (cutOff = FDR < = 0.05 & Log2FC > = 0) functions. Motif deviation scores were also plotted with ggplot.

## Data and code availability

Spatial mRNA-seq and spatial ATAC-seq data were deposited in the National Center for Biotechnology information Gene Expression Omibus (https://www.ncbi.nlm.nih.gov/geo/). The accession number will be available once the manuscript is accepted. We did not generate codes for novel algorithms. Custom codes were available upon request without restrictions.

## Authorship

Contributions: Y.L. performed bioinformatic analysis and wrote the manuscript; S.N., J.W., and L.M performed animal surgeries and collected tissues; S.M. provided technical support in epigenetic analysis; N.Z. analyzed the histology data; T.S. designed experiments and provided resources in animal surgery; J.F.D. developed hypotheses, designed experiments, and provided resources in animal surgery; Y.D. analyzed data and provided technical support in spatial ATAC-seq; Y.X. developed hypotheses, designed experiments, performed histology and spatial experiments, analyzed data, and wrote the manuscript.

## Acknowledgement

We thank Dr. Kam Leong for substantial scientific support in the initiation of this project. We thank Drs. Pengfei Guo, Parin Shah, Jennifer Garbarino, and Sebastian Werneburg for scientific discussion. Spatial mRNA sequencing was performed at the Human Immune Monitoring Core at Columbia University. We thank Tingting Wu, Xiaomei Wang, and Kevin Sun for the help with histology service (Nissl staining and bulk RNA extraction). High-resolution slide scanning of histology slides was performed at the Molecular Pathology Shared Resource of the Herbert Irving Comprehensive Cancer Center at Columbia University. We thank Jiaxin Zhu for assistance with computer resource access. We gratefully acknowledge the Allen Institute for Brain Science for the publicly available Allen Mouse Brain Atlas and *In Situ* Hybridization datasets, which facilitated the brain region annotation and analyses presented in this work. S.M. acknowledges support from the National Institutes of Health (NIH) through awards U01CA290442, R61CA297881, and R21CA301237, and the Mount Sinai Friedman Brain Institute Research Scholars Award. T.H.S. and J.M.W. acknowledge support from the National Institutes of Health (NIH) through awards R01NS142271 and R01NS120322. We acknowledge support from the Packard Fellowship for Science and Engineering (to Y.D.) and support from the Single Cell Spatial Analysis Program and the Department of Pathology at the University of Michigan (to Y.X.).

