## Supplements for "OmicGlaze: Spatial Multi-Omic Mapping of Traumatic Brain Injury"

### Supplementary Documents (8 Figures and 5 Tables)

S. Figure 1. Quality control and regional annotation for spatial transcriptomic samples.

S. Figure 2. *In situ* hybridization (ISH) images for marker genes.

S. Figure 3. Spatial gene modules of TBI_03 sample (Related to Figure 3).

S. Figure 4. Spatial gene modules of neuronal activities, stress and gliosis.

S. Figure 5. Spatial gene modules of activities related to thalamus, neuromodulation, hippocampus, neuroendocrine, striatum, and glial microenvironment.

S. Figure 6. Spatial gene modules of activities related to choroid plexus, fiber tracts, meninges, retrosplenial function, and hypothalamus.

S. Figure 7. Quality control metrics for spatial epigenetic samples.

S. Figure 8. Regional annotation for spatial epigenetic samples, marker motif volcano plot and *Mbp* gene expression.

S. Table 1. Gene set in each co-expressed gene module

S. Table 2. Quality control metrics for spatial transcriptomic dataset.

S. Table 3. Quality control metrics for spatial epigenetic dataset.

S. Table 4. Test statistical table using Wilcoxon rank sum test with continuity correction for module score distribution bar plot

S. Table 5. Gene set in Atf3 regulon

### Supplementary Figure 1

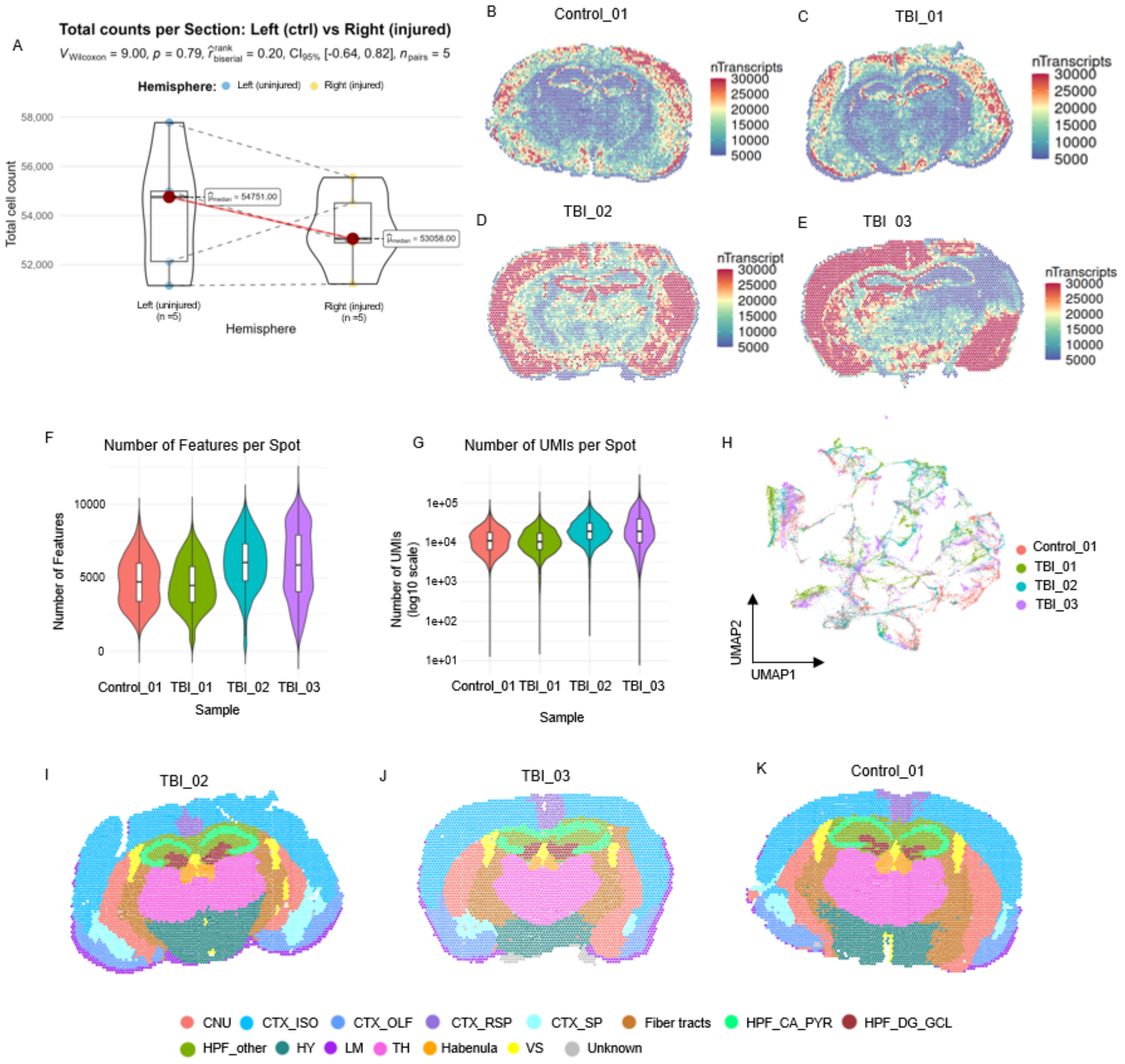


**Supplementary Figure 1. Quality control and regional annotation for spatial transcriptomic samples.**

**A** Comparison of total counts of cells from H&E staining image per section between the left (uninjured) and right (injured) hemispheres following traumatic brain injury. Each point is the summed total for a Section; colors denote hemisphere (Left ctrl vs Right injured). Lines connect the two hemispheres within the same Section; half-violins show the across- Section distribution. Paired Wilcoxon across Sections: V = 9.00, p = 0.79, $r_{(rank\text{-}biserial)}$ = 0.20 [95% CI −0.64, 0.82], n = 5. **B - E** Spatial distributions of the number of detected transcript per spot in A) control brain sample (Control_01) and TBI brain samples B) TBI_01, C) TBI_02, and D) TBI_03. **F** Violin plots of the number of detected features per spot across four samples. **G** Violin plots of the number of detected UMIs per spot across four samples. **H** UMAP projection of the integrated dataset from Control_01, TBI_01, TBI_02, and TBI_03 samples, colored by samples. **I - K** Spatial region annotation in K) Control_01, I) TBI_01, and J) TBI_02 samples, showing 14 anatomical regions. Abbreviations: CTX_ISO, isocortex; HPF_CA_PL, hippocampal formation cornu Ammonis pyramidal layer; TH, thalamus; HY, hypothalamus; LM, leptomeninges; HPF_other, rest of hippocampus region; CNU, cerebral nuclei; CTX_OLF, cortex olfactory areas; HPF_DG_GCL, hippocampal formation dentate gyrus granule layer; CTX_SP, cortical subplate; CTX_RSP, retrosplenial area; and VS, ventricular systems. All three figure use the same legend.

### Supplementary Figure 2

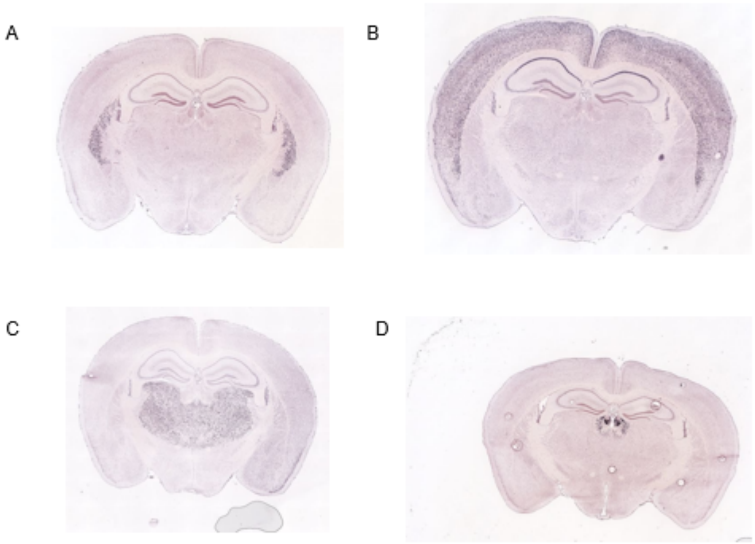


**Supplementary Figure 2. *In situ* hybridization images for marker genes.**

**A** *In situ* hybridization image for *Adora2a* expression in the adult mouse brain. Allen Mouse Brain Atlas [Internet]. 2006 Dec 06 [cited 2025 Oct 11]. Available from: https://mouse.brain-map.org/gene/show/11327. **B** *In situ* hybridization image for *Satb2* expression in the adult mouse brain. Allen Mouse Brain Atlas [Internet]. 2006 Dec 06 [cited 2025 Oct 11]. Available from: https://mouse.brain-map.org/gene/show/84457. **C** *In situ* hybridization image for *Synpo2* expression in the adult mouse brain. Allen Mouse Brain Atlas [Internet]. 2006 Dec 06 [cited 2025 Oct 11]. Available from: https://mouse.brain-map.org/gene/show/78699. **D** *In situ* hybridization image for *Gpr151* expression in the adult mouse brain. Allen Mouse Brain Atlas [Internet]. 2006 Dec 06 [cited 2025 Oct 11]. Available from: https://mouse.brain-map.org/gene/show/88694.

### Supplementary Figure 3

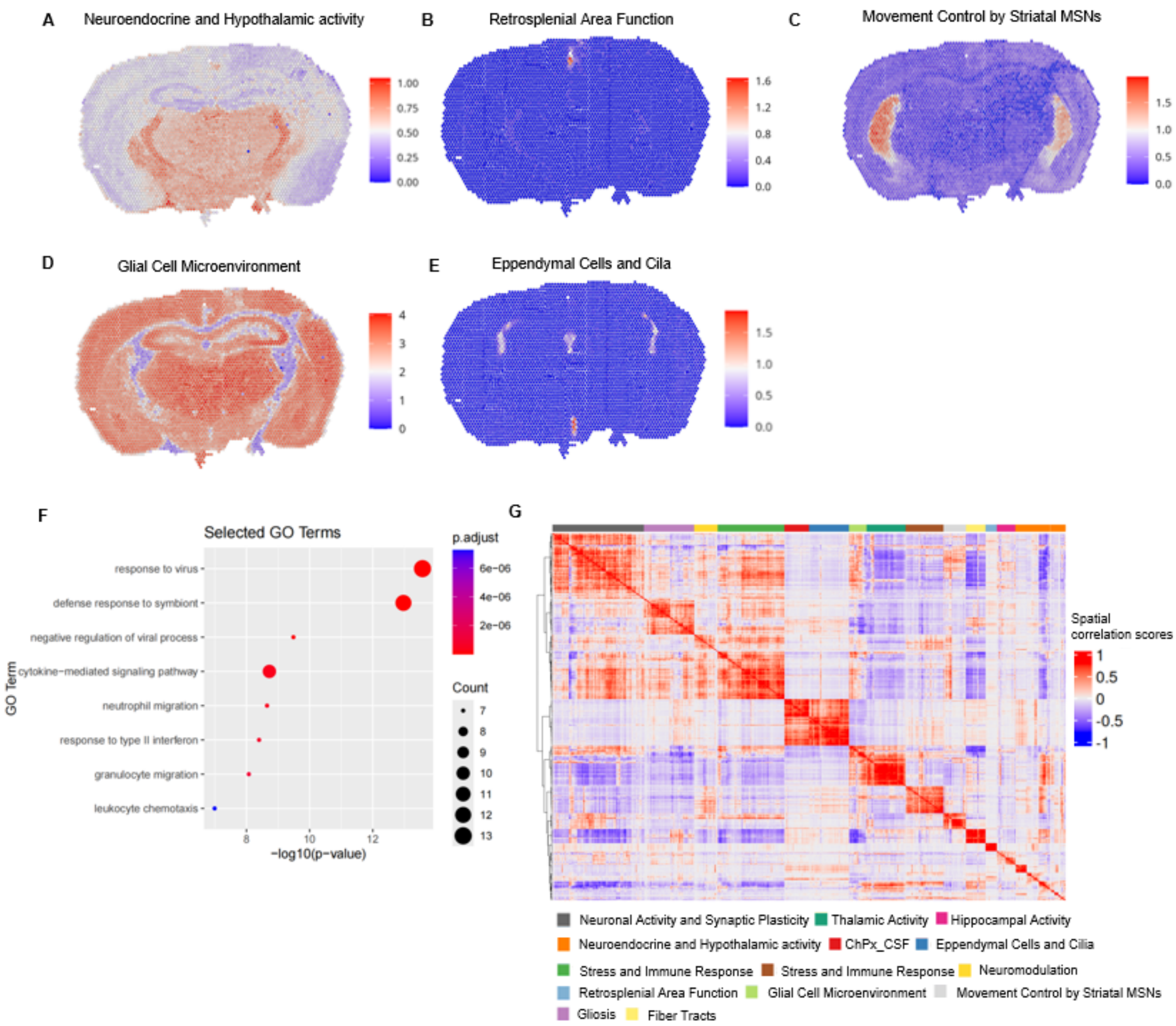


**Supplementary Figure 3. Spatial gene modules of TBI_03 sample (Related to Figure 3).**

**A - E** Spatial distribution of enrichment scores for gene modules (metagene clusters) A) *Neuroendocrine and Hypothalamic Activity*, B) *Retrosplenial Area Function*, C) *Movement Control by Striatal MSNs*, D) *Glial Cell Microenvironment*, E) *Eppendymal Cells and Cila* across brain tissue sections. A total of 14 gene modules were identified using Giotto and named according to their representative gene sets. Scores for nine modules are shown in main text Figure 3A. Each spot indicating the log-normalized average expression of all genes in the module; individual scale bars are used for each module. **F** Dot plot of manually selected top eight GO biological process terms significantly enriched in the gene set of the *Stress and Immune Response* module. **G** Heatmap showing the spatial correlation matrix of the 14 co-expressed gene modules identified, where red indicates positive correlations and blue indicates negative correlations. Genes are grouped by spatial co-expression modules (color bar on top).

### Supplementary Figure 4

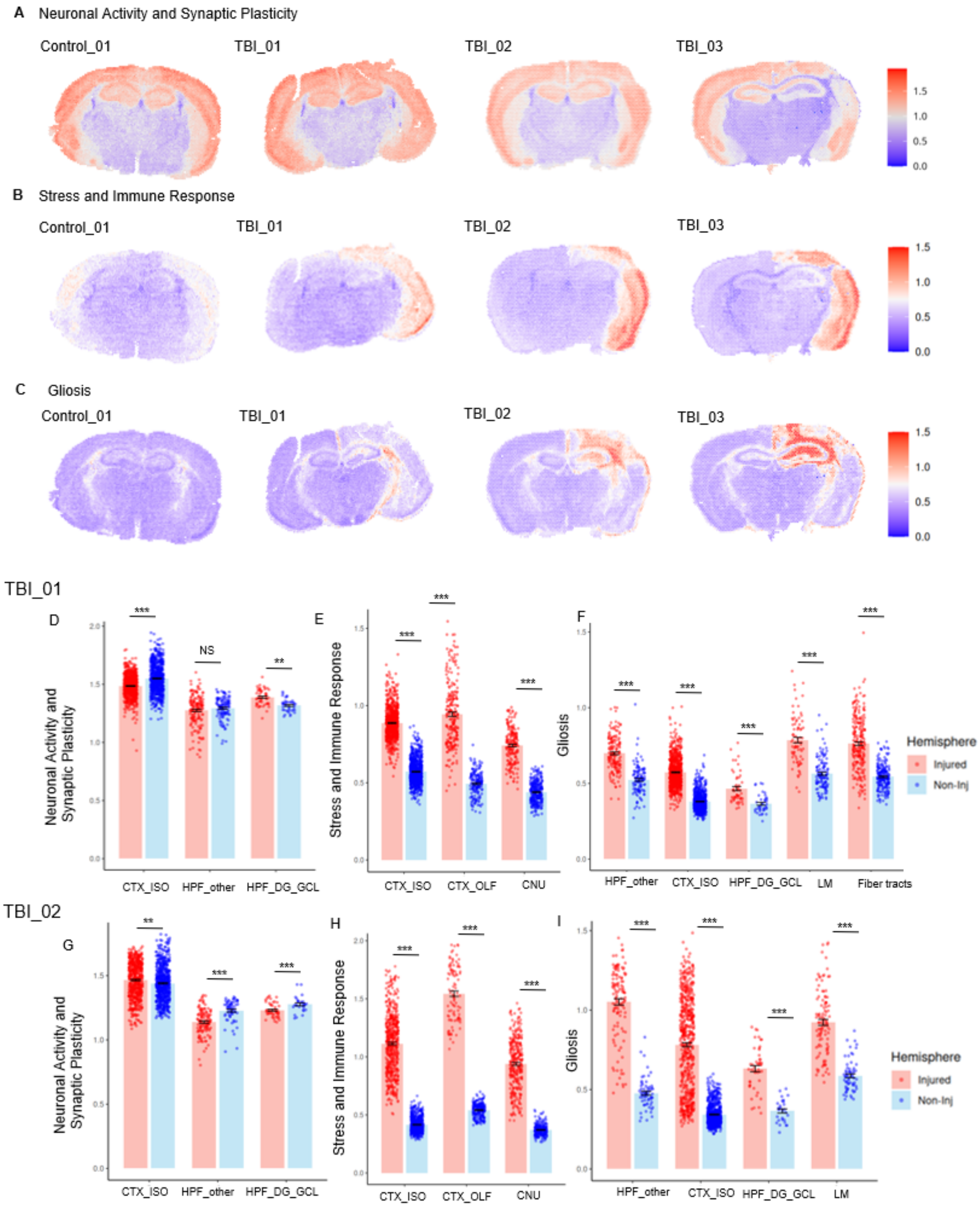


**Supplementary Figure 4. Spatial gene modules of neuronal activities, stress and gliosis.**

**A - C** Spatial distribution of enrichment scores for the three highlight gene modules for all four samples: Control_01, TBI_01, TBI_02 and TBI_03 samples. A) *Neuronal Activity and Synaptic Plasticity*, B) *Stress and Immune Response*, C) *Gliosis.* Same scale bar is used for each module. **D - F** Bar plots illustrating the distribution of module scores for D) the *Neuronal Activity and Synaptic Plasticity*, E) *Stress and Immune Response*, and F) *Gliosis* modules at each spot for selected paired regions on the injured (red) and uninjured (blue) sides of the TBI_01 brain. Each plot includes three or five regions, with each dot representing a single spot from the indicated region. Wilcoxon rank-sum test p-values for each comparison are shown on the plots. **G - I** Bar plots illustrating the distribution of module scores for D) the *Neuronal Activity and Synaptic Plasticity*, E) *Stress and Immune Response*, and F) *Gliosis* modules at each spot for selected paired regions on the injured (red) and uninjured (blue) sides of the TBI_02 brain. Each plot includes three or four regions, with each dot representing a single spot from the indicated region. Wilcoxon rank-sum test p-values for each comparison are shown on the plots.

### Supplementary Figure 5

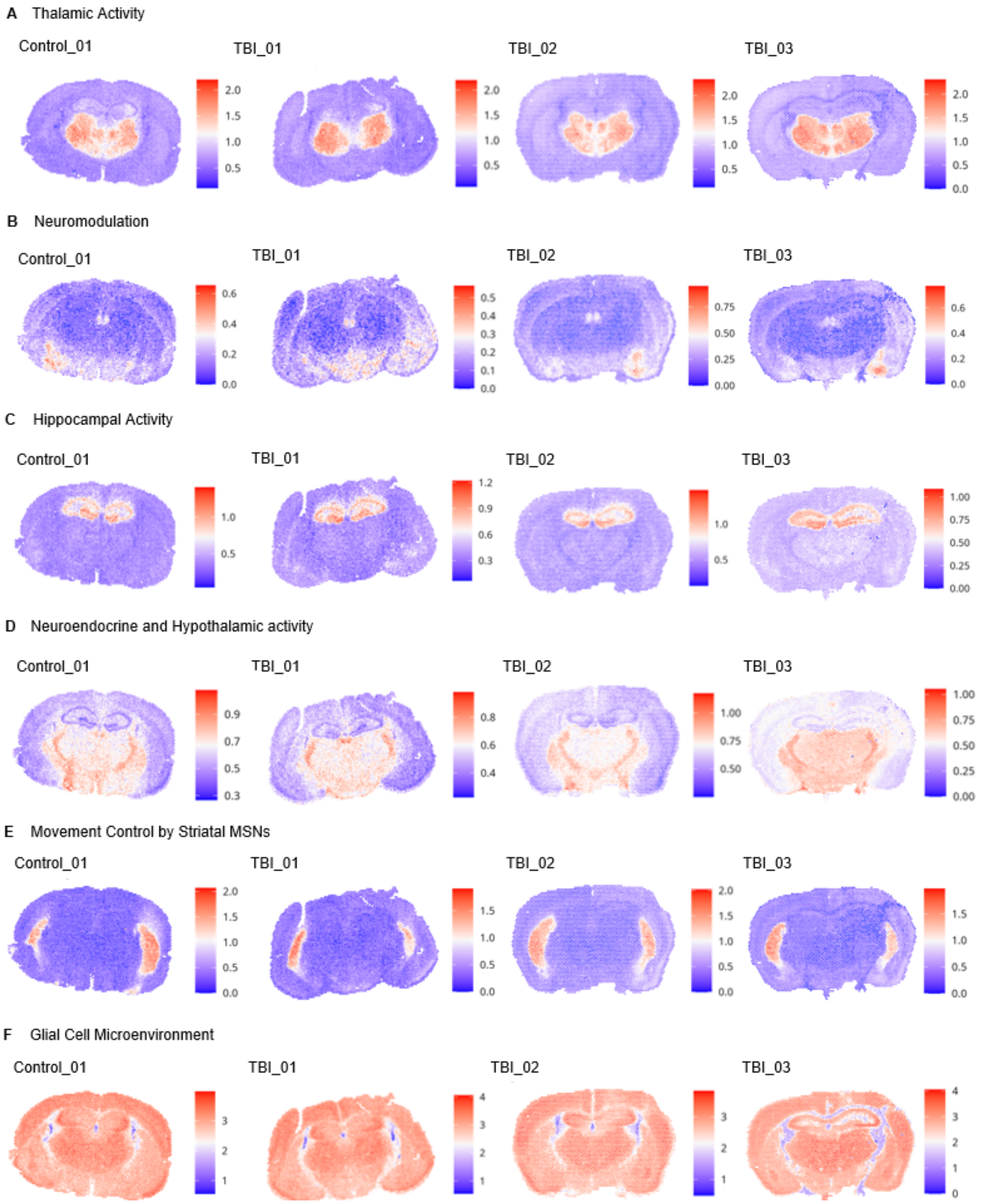


**Supplementary Figure 5. Spatial gene modules of activities related to thalamus, neuromodulation, hippocampus, neuroendocrine, striatum, and glial microenvironment.**

**A - F** Spatial distribution of enrichment scores for the first six gene modules identified earlier across all four brain tissue sections. Each spot indicating the log-normalized average expression of all genes in the module; individual scale bars are used for each sample and each module.

### Supplementary Figure 6

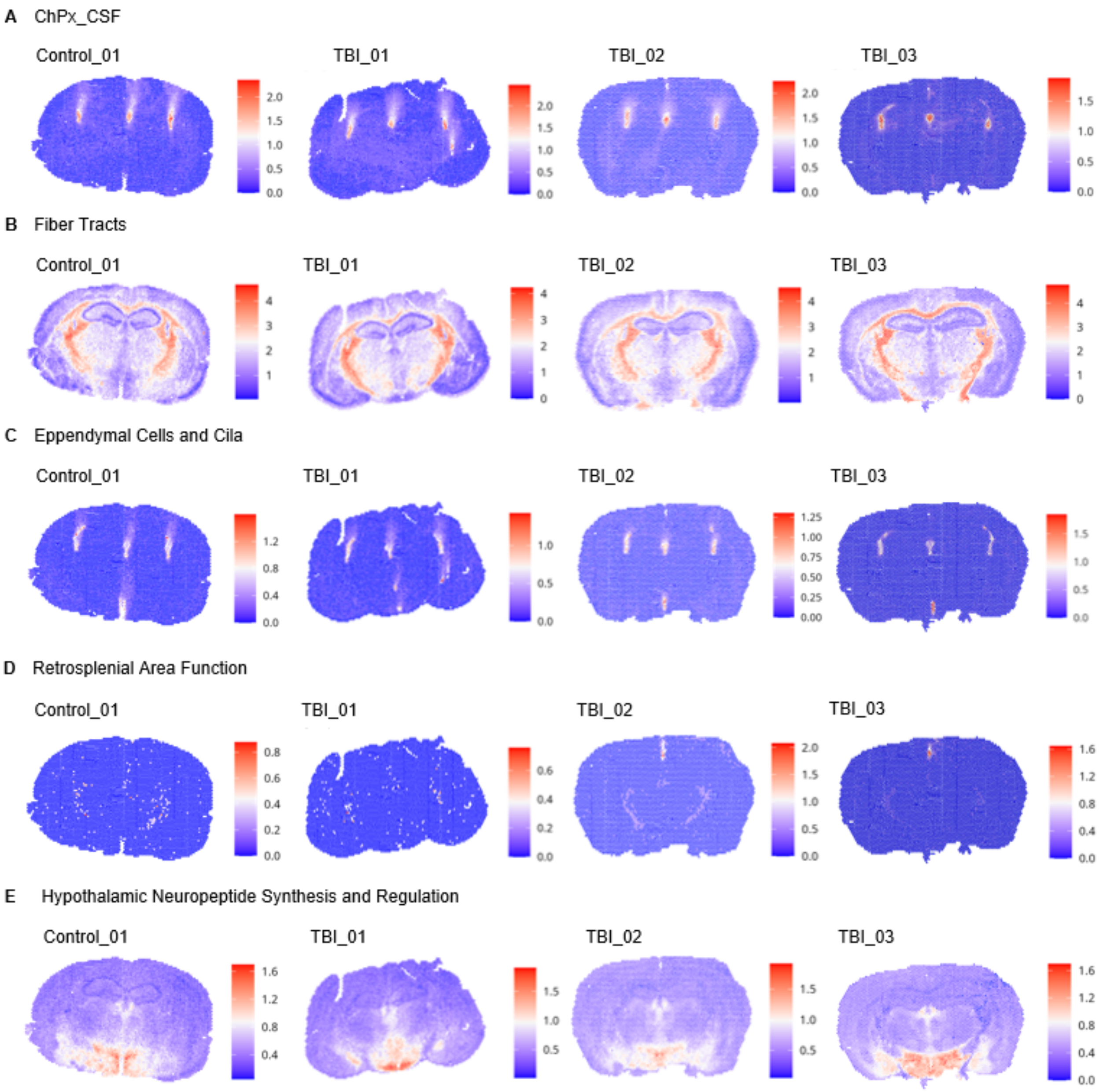


**Supplementary Figure 6. Spatial gene modules of activities related to choroid plexus, fiber tracts, meninges, retrosplenial function, and hypothalamus.**

**A - E** Spatial distribution of enrichment scores for the rest five gene modules identified earlier across all four brain tissue sections. Each spot indicating the log-normalized average expression of all genes in the module; individual scale bars are used for each sample and each module.

### Supplementary Figure 7

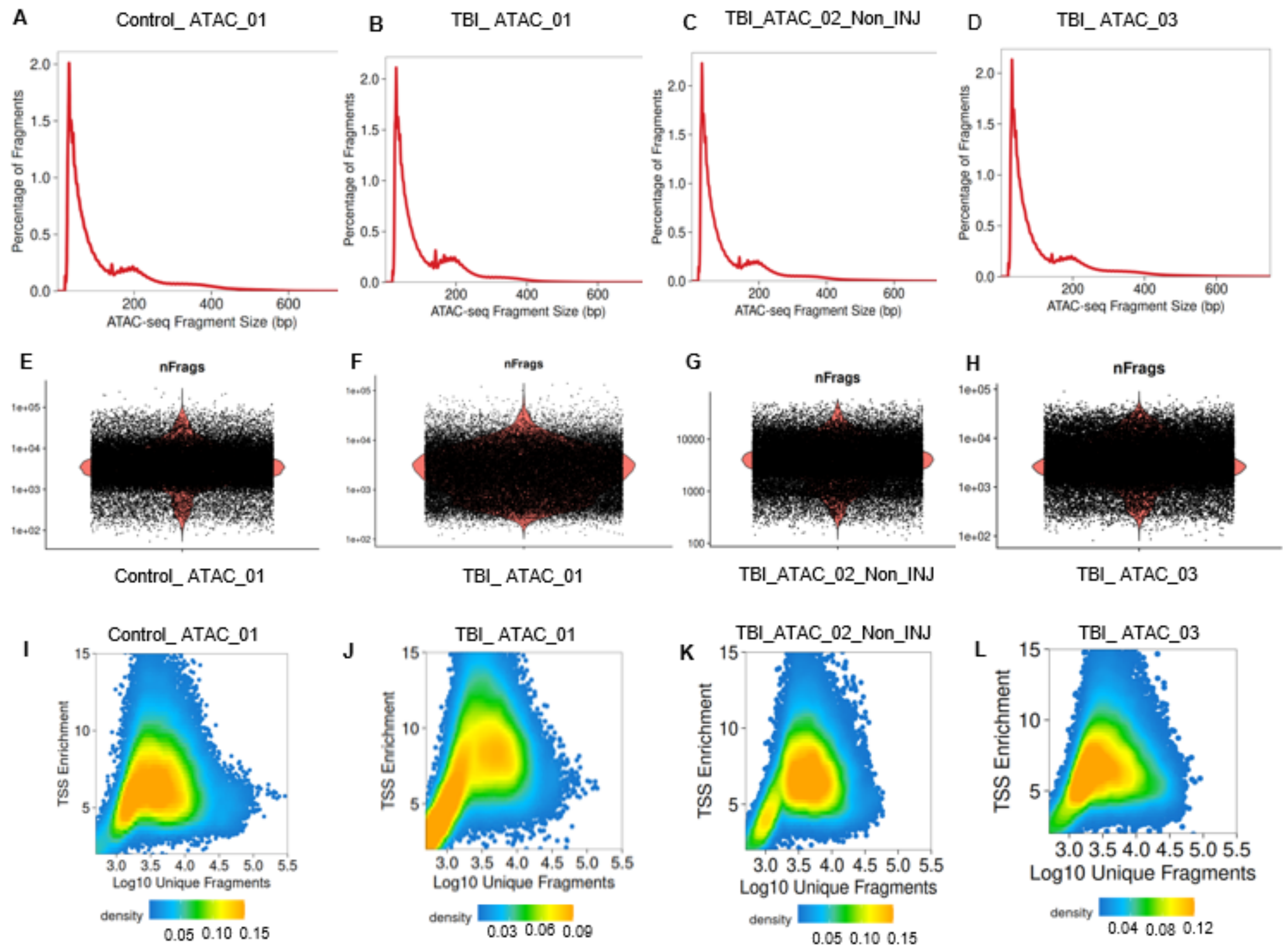


**Supplementary Figure 7. Quality control metrics for spatial epigenetic samples.**

**A - D** Distributions of spatial ATAC-seq fragment sizes for A) Control_ ATAC_01, B) TBI_ ATAC_01, C) TBI_ATAC_02_Non_INJ, and D) TBI_ ATAC_03, represented as the percentage of total fragments (y-axis) against fragment size in base pairs (x-axis). **E - H** Violin plots of distributions of fragment counts (nFrags) across individual spots for E) Control_ ATAC_01, F) TBI_ ATAC_01, G) TBI_ATAC_02_Non_INJ, and H) TBI_ ATAC_03. Each dot represents a spot in the chip, with density shown on the sides. **I - L** Scatter density plot showing transcription start site (TSS) enrichment (y-axis) versus the number of unique fragments per cell (x-axis, log10 scale) for I) Control_ ATAC_01, J) TBI_ ATAC_01, K) TBI_ATAC_02_Non_INJ, and L) TBI_ ATAC_03.

### Supplementary Figure 8

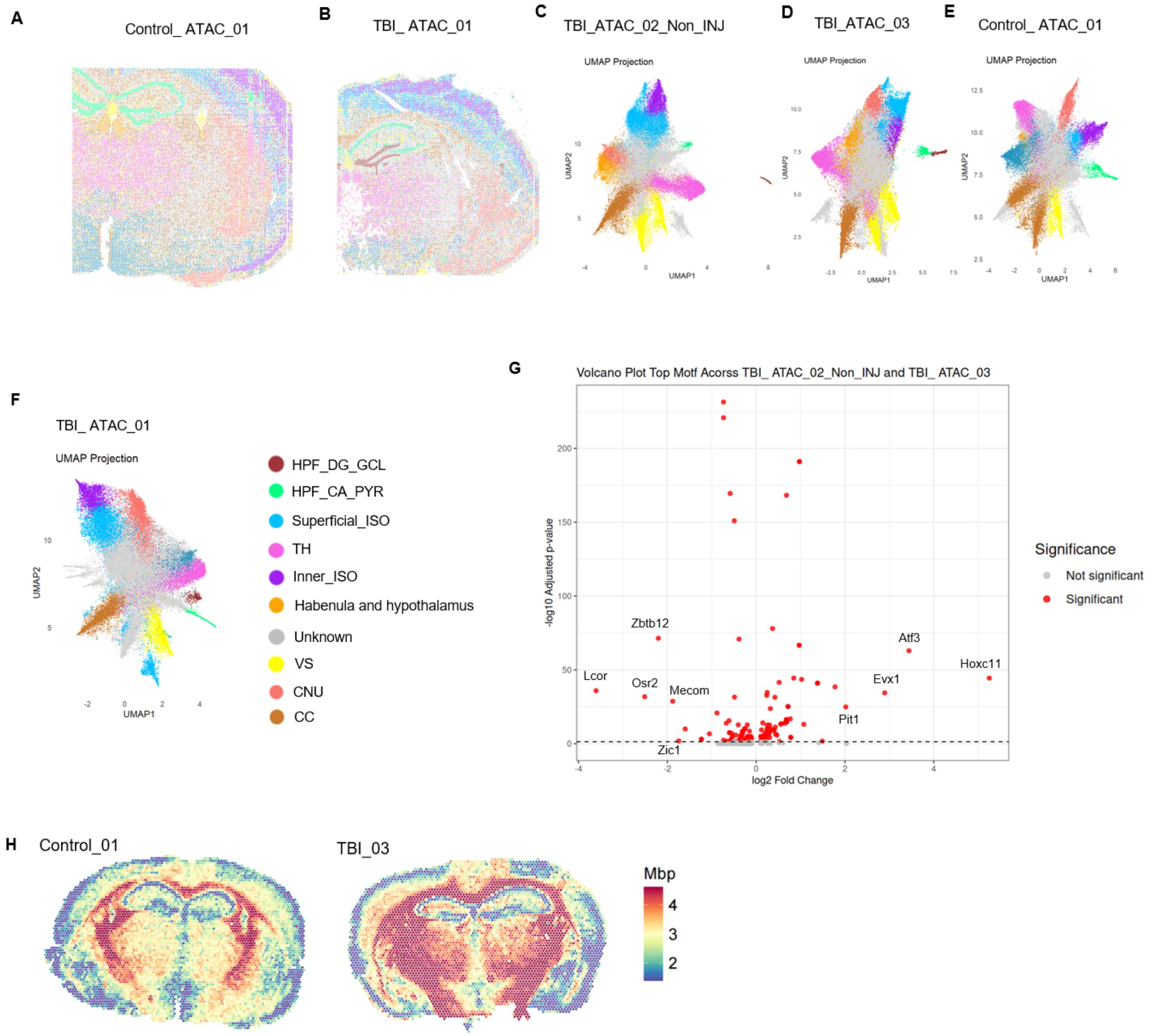


**Supplementary Figure 8. Regional annotation for spatial epigenetic samples, marker motif volcano plot and *Mbp* gene expression.**

**A - B** Spatial distribution of clusters in ATAC-seq data mapping for A) Control_ATAC_01 and B) TBI_ATAC_01. **E – F** UMAP projection colored by clusters for each sample. Significant clusters are highlighted with abbreviations: HPF_CA_PYR, hippocampal formation cornu ammonis pyramidal layer, HPF_DG_GCL, hippocampal formation dentate gyrus granule cell layer, Inner_ISO, inner layer of isocortex, Superfical_ISO, superficial layer of isocortex, CNU, cerebral nuclei, TH, thalamus, CC, Corpus callosum, VS, Ventricle system. **G** Volcano plot showing differential motif accessibility between TBI_ATAC_02_Non_INJ and TBI_ATAC_03 samples. Each red dot represents a transcription factor motif, with the x-axis showing $\log_2$ fold change and the y-axis indicating $-\log_{10}$ adjusted p-value. The dashed line marks the significance threshold. **H** Spatial mapping of *Mbp* gene in Control_01 and TBI_03 samples.

**Supplementary Table 1. Genes in each co-expressed gene module**

| Module Name | Genes in the module |
|---|---|
| 1_Neuronal Activity_Synaptic Plasticity | 2010300C02Rik, Adra1d, Aldoc, Arpp19, Arpp21, Atp1a1, Baiap2, Basp1, Camk2a, Camk2n1, Cck, Ccn2, Ccn3, Cfl1, Chn1, Cnksr2, Cplx3, Ctxn1, Cyp26b1, Ddn, Dkk3, Doc2a, Enc1, Epop, Esrp2, Fam81a, Fezf2, Fhl2, Gas7, Gda, Glt8d2, Gm765, Gprin1, Homer1, Hpca, Hpcal4, Hs3st2, Icam5, Igfbp6, Igfn1, Inka2, Itpka, Kcnh4, Kcnip2, Lamp5, Lingo1, Lmo4, Ly6h, Mef2c, Mmp17, Myl4, Nap1l5, Ndrg2, Neurod2, Neurod6, Ngef, Nptxr, Nr4a2, Nrgn, Nrn1, Nxph3, Olfm1, Ovol2, Pcsk1n, Pde1a, Pde2a, Pdzrn3, Ppp3ca, Psd, Robo3, Rprm, Rspo2, Rsrp1, Rtn4r, Rtn4rl2, Satb2, Slc17a7, Slc1a2, Slc22a17, Slc30a3, Snca, Sv2b, Synpo, Tafa1, Tmem215, Trbc2, Urah, Vxn, Wnt10a |
| 2_Thalamic Activity | AI593442, Abhd12b, Adamts19, Adarb1, Adgrb1, Amotl1, Atp2a1, Ccdc136, Cit, Dbndd1, Epn3, Eps8l2, Gabrd, Gnal, Hspa12a, Kcnc2, Mrvi1, Nexn, Ntng1, Ogfrl1, Patj, Pcp4, Plcb4, Plekhg1, Prkcd, Ptpn3, Ptpn4, Rab37, Ramp3, Rasd1, Rora, Rorb, Scn1a, Slitrk6, Synpo2, Tcf7l2, Tnnt1, Vav3, Zic1 |
| 3_Hippocampal Activity | C1ql2, Cabp7, Ccbe1, Crlf1, Dsp, Fibcd1, Klk8, Lct, Lypd8, Ntf3, Olfr61, Rtl3, Selenow, Shisa6, Slc9a4, Smoc2, Spink8 |
| 4_Neuroendocrine and Hypothalamic activity | Actg2, Agt, Aplp2, App, Atp6v1g3, Avp, Cdh3, Chat, Chrna3, Etnppl, Frem3, Gad2, Gbx2, Gh, Gm5741, Gpr151, Htr5b, Igf2, Irx2, Isl1, Itih3, Lhx8, Mpz, Myoc, Nmbr, Nppa, Osbpl1a, Oxt, Paip2, Pln, Pomc, Pou4f1, Prl, Ptprq, Pvalb, Six3, Slc13a4, Slc18a3, Slc26a7, Slc5a7, Slc6a11, Snx31, Sparc, Stmn4, Tmem182, Tmod2, Tpsab1, Tpsb2, Wfikkn2 |

| | |
|---|---|
| 5_ChPx_CSF | Aqp1, Calml4, Cldn2, Clic6, Defb11, Defb9, Ecrg4, F5, Folr1, Kcne2, Kcnj13, Kl, Lmx1a, Mfrp, Oca2, Prr32, Slc16a8, Slc4a5, Sostdc1, Sult1c2, Tmem72, Tmprss11a, Ttr, Wdr86 |
| 6_Eppendymal Cells and Cilia | 1700001C02Rik, 1700012B09Rik, 1700024G13Rik, 2410004P03Rik, Acox2, Ak7, Ak9, Anxa8, Asb14, Capsl, Ccdc153, Ccdc162, Ccdc180, Cfap299, Cfap43, Cfap52, Cfap65, Col8a2, Dnah1, Dnah12, Dnali1, Drc7, Dynlrb2, Fam216b, Lbp, Lrrc23, Lrrc36, Lrrc74b, Mia, Olfr1507, Pifo, Prr29, Rsph1, Sntn, Spag16, Stoml3, Tctex1d4, Tmem212, Unc5cl |
| 7_Stress_Immune Response | Abhd2,Acan,Alkal2,Anp32a,Arc,Bdnf,Car12,Cdkn1a,Clec18a,Crhbp,Cryba2,Drd4,Dusp4,Dusp5,Efhd2,Egr1,Egr2,Egr3,Egr4,Fam83g,Fndc9,Fos,Fosb,Gadd45b,Gal,Hsbp1,Hspb1,Ifi202b,Ifrd1,Igsf9b,Inhba,Jsrp1,Junb,Klhl40,Krt16,Krt6a,Lif,Ly6g6c,Ncs1,Npas4,Nptx2,Npw,Nr4a1,Pfn2,Pgm2l1,Plcb1,Prkar1b,Ptgs2,Ptp4a3,Rtn1,S100a10,Scamp1,Schip1,Sdc1,Serpinb2,Serpine1,Sh3bp5,Slc6a5,Socs3,Spred1,Sstr2,Stx1b,Tll1,Tpm1,Vgf |
| 8_Hypothalamic Neuropeptide Synthesis and Regulation | Abhd2, Acan, Alkal2, Anp32a, Arc, Bdnf, Car12, Cdkn1a, Clec18a, Crhbp, Cryba2, Drd4, Dusp4, Dusp5, Efhd2, Egr1, Egr2, Egr3, Egr4, Fam83g, Fndc9, Fos, Fosb, Gadd45b, Gal, Hsbp1, Hspb1, Ifi202b, Ifrd1, Igsf9b, Inhba, Jsrp1, Junb, Klhl40, Krt16, Krt6a, Lif, Ly6g6c, Ncs1, Npas4, Nptx2, Npw, Nr4a1, Pfn2, Pgm2l1, Plcb1, Prkar1b, Ptgs2, Ptp4a3, Rtn1, S100a10, Scamp1, Schip1, Sdc1, Serpinb2, Serpine1, Sh3bp5, Slc6a5, Socs3, Spred1, Sstr2, Stx1b, Tll1, Tpm1, Vgf |
| 9_Neuromodulation | 6430628N08Rik, Adcyap1, Ahi1, Atp2b4, Barhl2, Ccdc42, F2rl2, Gjb3, Hmcn2, Krt75, Mep1a, Nts, Nxph4, Pitx2, Pth2, Rrad, Scn5a, Slc6a20b, Styk1, Tac2, Tprg, Ucn3, Uox |

| | |
|---|---|
| 10_Retrosplenial Area Function | Crx, Gm45716, Gnat2, Gnb3, Lhx4, Lrit1, Pde6c, Rbp3, Spink4, Tph1 |
| 11_Glial Cell Microenvironment | Atp1b1, Bsn, Calm1, Camk2b, Dynll2, Eno2, Gpm6a, Grin1, Rgs4, Sept5, Snap25, Snhg11, Stmn3, Stxbp1, Sult4a1, Syp, Uchl1 |
| 12_Movement Control by Striatal MSNs | Actn2, Adora2a, Chia1, Crabp1, Drd2, Gpr6, Gpr88, Ido1, Krt9, Lrrc10b, Pde10a, Penk, Ppp1r1b, Rem2, Rgs9, Rxrg, Scn4b, Sh3rf2, Syndig1l, Tac1, Tpbg, Wfs1 |
| 13_Gliosis | 1700022I11Rik, A2m, Adam8, Atf3, Bcas1, Bst2, C4b, Ccl11, Ccl5, Ccl7, Ccn1, Cd44, Cxcl10, Flnc, Fth1, Gbp2, Gfap, Gm614, Hist1h2bk, Hist1h3b, Hist1h3c, Hmox1, Ifit1, Ifit3, Ifitm3, Igfbp4, Il1rn, Il6, Irf7, Lcn2, Lgals3, Mt1, Mt2, Oasl1, Oasl2, Ptx3, Rapgef4, Rsad2, Rtp4, S100a4, S100a6, Serpina3n, Steap4, Tgm1, Thbs4, Tm4sf1, Tmsb4x, Tubb6, Usp18, Vim |
| 14_Fiber Tracts | Cldn11, Cnp, Ermn, Fa2h, Gatm, Gpr37, Mag, Mal, Mbp, Mobp, Mog, Plekhb1, Plp1, Ppp1r14a, Ptgds, Scd2, Sept4, Trf, Ugt8a |

**Supplementary Table 2. Quality control metrics for spatial transcriptomic dataset**.

| **Sample** | **Control_01** | **TBI_01** | **TBI_02** | **TBI_03** |
|---|---|---|---|---|
| Original Identity | YXV01 | YXV02 | YXV03 | YXV04 |
| Injury Model | No injury | Fluid Percussion Injury | Controlled Cortical Impact | Controlled Cortical Impact |
| Injury Plane | Not Applicable | Peri-injury | Peri-injury | Injury Center |
| Post-TBI (Hours) | Not applicable | 3 | 24 | 24 |
| Fixation | Unfixed | Unfixed | Unfixed | Unfixed |
| Tissue Preservation | Fresh Frozen | Fresh Frozen | Fresh Frozen | Fresh Frozen |
| Tissue Thickness (microns) | 10 | 10 | 10 | 10 |
| Number of Spots Under Tissue | 5,840 | 5,627 | 4,704 | 4,842 |
| Mean Reads per Spot | 37,628 | 45,701 | 54,191 | 60,984 |
| Median Genes per Spot | 4,720 | 4,479 | 6,038 | 5,858 |
| Median UMI Counts per Spot | 10,988 | 10,429 | 19,035 | 19,038 |
| Total Genes Detected | 16,829 | 16,847 | 17,364 | 17,465 |
| Total Number of Reads | 219,748,014 | 257,156,749 | 254,913,522 | 295,283,362 |
| Fraction Reads in Spots Under Tissue | 93.9% | 91.4% | 98% | 97.8% |
| Sequencing Saturation | 58.3% | 66.4% | 51.2% | 46.3% |

**Supplementary Table 3. Quality control metrics for spatial epigenetic dataset**.

| **Sample** | **Control_ ATAC_01** | **TBI_ ATAC_01** | **TBI_ ATAC_02_Non_INJ** | **TBI_ ATAC_03** |
|---|---|---|---|---|
| Original Ident | FPI_CTRL_73 | FPI_AD13_73 | CCI_Lateral_68 | CCI_Injury_67 |
| Injury Model | No injury | Fluid Percussion Injury | Controlled Cortical Impact | Controlled Cortical Impact |
| Post-TBI (Hours) | Not applicable | 3 | 24 | 24 |
| Brain Hemisphere | Non_INJ | INJ | Non_INJ | INJ |
| TSS enrichment score | 5.14 | 6.12 | 5.66 | 5.28 |
| FRIP (Fraction of Reads in Peaks) | 0.28 | 0.34 | 0.29 | 0.29 |
| Median Fragements per spot | 3700 | 2600 | 4300 | 3200 |
| Sequenced read pairs | 961 million | 911 million | 687 million | 674 million |
| Fraction Duplicates | 0.39 | 0.65 | 0.27 | 0.32 |

**Supplementary Table 4. Test statistical table using Wilcoxon rank sum test with continuity correction for module score distribution bar plot**

| Module | Region | Comparsion | N1 | N2 | p.value |
|---|---|---|---|---|---|
| Neuronal Activity and Synaptic Plasticity | CTX_ISO | CTRL vs TBI_Un_Inj | 1295 | 571 | 1.046778e-19 |
| Neuronal Activity and Synaptic Plasticity | HPF_other | CTRL vs TBI_Un_Inj | 369 | 74 | 2.410478e-13 |
| Neuronal Activity and Synaptic Plasticity | HPF_DG_GCL | CTRL vs TBI_Un_Inj | 74 | 39 | 8.115700e-04 |
| Neuronal Activity and Synaptic Plasticity | CTX_ISO | TBI_Un_Inj vs TBI_Inj | 571 | 561 | 5.175698e-127 |
| Neuronal Activity and Synaptic Plasticity | HPF_other | TBI_Un_Inj vs TBI_Inj | 74 | 163 | 3.590746e-27 |
| Neuronal Activity and Synaptic Plasticity | HPF_DG_GCL | TBI_Un_Inj vs TBI_Inj | 39 | 51 | 5.795168e-16 |
| Neuronal Activity and Synaptic Plasticity | CTX_ISO | CTRL vs TBI_Inj | 1295 | 561 | 3.292638e-201 |
| Neuronal Activity and Synaptic Plasticity | HPF_other | CTRL vs TBI_Inj | 369 | 163 | 1.106615e-71 |
| Neuronal Activity and Synaptic Plasticity | HPF_DG_GCL | CTRL vs TBI_Inj | 74 | 51 | 2.622281e-21 |
| Stress and Immune Response | CTX_ISO | CTRL vs TBI_Un_Inj | 1295 | 571 | 7.080100e-241 |
| Stress and Immune Response | CTX_OLF | CTRL vs TBI_Un_Inj | 325 | 199 | 1.092568e-81 |
| Stress and Immune Response | CNU | CTRL vs TBI_Un_Inj | 731 | 162 | 2.949801e-87 |
| Stress and Immune Response | CTX_ISO | TBI_Un_Inj vs TBI_Inj | 571 | 561 | 3.533880e-184 |
| Stress and Immune Response | CTX_OLF | TBI_Un_Inj vs TBI_Inj | 199 | 96 | 5.203211e-44 |

| | | | | | |
|---|---|---|---|---|---|
| Stress and Immune Response | CNU | TBI_Un_Inj vs TBI_Inj | 162 | 253 | 3.149149e-66 |
| Stress and Immune Response | CTX_ISO | CTRL vs TBI_Inj | 1295 | 561 | 8.129480e-218 |
| Stress and Immune Response | CTX_OLF | CTRL vs TBI_Inj | 325 | 96 | 3.686882e-50 |
| Stress and Immune Response | CNU | CTRL vs TBI_Inj | 731 | 253 | 9.624481e-117 |
| Gliosis | CTX_ISO | CTRL vs TBI_Un_Inj | 1295 | 571 | 1.838413e-43 |
| Gliosis | HPF_other | CTRL vs TBI_Un_Inj | 369 | 74 | 1.067733e-13 |
| Gliosis | HPF_DG_GCL | CTRL vs TBI_Un_Inj | 74 | 39 | 1.993655e-01 |
| Gliosis | LM | CTRL vs TBI_Un_Inj | 71 | 90 | 2.982193e-23 |
| Gliosis | CTX_ISO | TBI_Un_Inj vs TBI_Inj | 571 | 561 | 1.679720e-148 |
| Gliosis | HPF_other | TBI_Un_Inj vs TBI_Inj | 74 | 163 | 1.218463e-33 |
| Gliosis | HPF_DG_GCL | TBI_Un_Inj vs TBI_Inj | 39 | 51 | 1.789483e-15 |
| Gliosis | LM | TBI_Un_Inj vs TBI_Inj | 90 | 60 | 4.074248e-22 |
| Gliosis | CTX_ISO | CTRL vs TBI_Inj | 1295 | 561 | 3.732777e-167 |
| Gliosis | HPF_other | CTRL vs TBI_Inj | 369 | 163 | 6.249293e-75 |
| Gliosis | HPF_DG_GCL | CTRL vs TBI_Inj | 74 | 51 | 2.622281e-21 |
| Gliosis | LM | CTRL vs TBI_Inj | 71 | 60 | 7.775965e-23 |

**Supplementary Table 5. Gene set in Atf3 regulon**

| Gene Name | Normalized Enrichment Score | Gene Name | Normalized Enrichment Score |
|---|---|---|---|
| Adamts9 | 7.32 | Zmiz1 | 7.32 |
| Ahnak | 7.32 | C3ar1 | 3.78 |
| Anxa2 | 7.32 | Cd14 | 3.78 |
| Bcl3 | 7.32 | Cd68 | 3.78 |
| Cd44 | 7.32 | Clic1 | 3.78 |
| Cdh5 | 7.32 | Cryba2 | 3.78 |
| Csrnp1 | 7.32 | Dmkn | 3.78 |
| Efhd2 | 7.32 | Dusp5 | 3.78 |
| Emp1 | 7.32 | Ecscr | 3.78 |
| Emp3 | 7.32 | Fos | 3.78 |
| Flnc | 7.32 | Ifit3 | 3.78 |
| Fosl1 | 7.32 | Sphk1 | 3.78 |
| Hbegf | 7.32 | Ccl12 | 3.27 |
| Hmox1 | 7.32 | Ccl3 | 3.27 |
| Hs3st2 | 7.32 | Egr2 | 3.27 |
| Hspb1 | 7.32 | Ifrd1 | 3.27 |
| Ier3 | 7.32 | Stat3 | 3.27 |
| Igsf9b | 7.32 | Arl4d | 3.05 |
| Il11 | 7.32 | Atf3 | 3.05 |
| Itga5 | 7.32 | Btg2 | 3.05 |

| | | | |
|---|---|---|---|
| Itpkc | 7.32 | Ch25h | 3.05 |
| Junb | 7.32 | Epha2 | 3.05 |
| Kdm6b | 7.32 | Fosb | 3.05 |
| Lgals3 | 7.32 | Gem | 3.05 |
| Lif | 7.32 | Gpr3 | 3.05 |
| Msn | 7.32 | Ier2 | 3.05 |
| Myh9 | 7.32 | Inhba | 3.05 |
| Npas4 | 7.32 | Jun | 3.05 |
| Pdlim4 | 7.32 | Klf6 | 3.05 |
| Prss23 | 7.32 | Maff | 3.05 |
| Rara | 7.32 | Ppp1r15a | 3.05 |
| Rgs2 | 7.32 | Relb | 3.05 |
| Rrad | 7.32 | Rhoj | 3.05 |
| S100a10 | 7.32 | Sik1 | 3.05 |
| Serpine1 | 7.32 | Tnfaip3 | 3.05 |
| Sertad1 | 7.32 | Trib1 | 3.05 |
| Sh3pxd2b | 7.32 | Ccdc80 | 3.03 |
| Slc6a8 | 7.32 | Dab2 | 3.03 |
| Srxn1 | 7.32 | Nckap1l | 3.03 |
| Stk40 | 7.32 | Nedd9 | 3.03 |
| Tagln2 | 7.32 | Nfkbiz | 3.03 |
| Tgfbi | 7.32 | Pik3ap1 | 3.03 |
| Tnfrsf12a | 7.32 | Rasip1 | 3.03 |

| | | | |
|---|---|---|---|
| Vim | 7.32 | Tgif1 | 3.03 |
| Zfp36 | 7.32 | | |
